# Sex-dependent interactions between prodromal intestinal inflammation and LRRK2 G2019S in mice promote symptoms of Parkinson’s disease

**DOI:** 10.1101/2022.11.30.518591

**Authors:** Ping Fang, Hannah Espey, Lewis W. Yu, Gulistan Agirman, Kai Li, Yongning Deng, Jamie Lee, Haley Hrncir, Arthur P. Arnold, Elaine Y. Hsiao

## Abstract

Gastrointestinal (GI) disruptions, such as inflammatory bowel disease (IBD), are common prodromal symptoms of Parkinson’s disease (PD), but how they may impact risk for PD remains poorly understood. Herein, we provide evidence that prodromal intestinal inflammation expedites and exacerbates PD symptoms in rodent carriers of the human PD risk allele LRRK2 G2019S in a sex-dependent manner. Chronic intestinal damage in genetically predisposed male, but not female, mice promotes α-synuclein aggregation in the substantia nigra, elevated α-synuclein loads in microglia, loss of dopaminergic neurons, and motor impairment. This male bias in gene-environment interaction is preserved in gonadectomized males, and similarly conferred by sex chromosomal complement in gonadal females expressing human LRRK2 G2019S, revealing that XY chromosomes, not testicular hormones, mediate the male bias in the gut-brain-driven risk for endophenotypes of PD. The early onset and heightened severity of neuropathological and behavioral outcomes in male LRRK2 G2019S mice is preceded by the accumulation of α-synuclein in the colon, increases in α-synuclein-positive macrophages in the colonic lamina propria, and elevations in α-synuclein loads within microglia in the substantia nigra. Taken together, these data reveal that prodromal intestinal inflammation promotes the pathogenesis of PD symptoms in male carriers of LRRK2 G2019S, through mechanisms that depend on genotypic sex and involve early accumulation of α-synuclein in myeloid cells within the gut and brain.

---

Parkinson’s disease (**PD**) is an aging-related neurodegenerative disorder that is rising in prevalence. The global burden of PD has more than doubled since 1990 to an estimated 6.2 million individuals in 2015 (*1*), with a projected 14 million people afflicted with PD by 2040 (*2*). While currently available therapies offer symptomatic relief, the progressive nature of PD leads to drug-resistant motor impairments (*3*). The average age of PD diagnosis is 60 years, wherein 50-60% of dopaminergic neurons have already been lost (*3*). As such, the disease process begins well before diagnostic symptoms of PD arise, and there is a pressing need to understand early causes of PD toward enabling early detection and intervention.

Gastrointestinal dysfunction and inflammation are increasingly implicated in PD. Subsets of PD patients experience recurrent constipation, dysphagia, and small intestinal bacterial overgrowth, among other gastrointestinal disturbances, which can precede the onset of motor symptoms by several years (*4-6*). PD is further linked to intestinal inflammation as its incidence is over 20% higher in individuals with inflammatory bowel disease (**IBD**) relative to non-IBD controls (*7, 8*). The occurrence of PD among IBD patients is decreased in those who previously received anti-tumor necrosis factor therapy (*9*), suggesting that peripheral immunosuppression of intestinal inflammation reduces risk for PD. Moreover, increasing evidence supports Braak’s hypothesis that synucleinopathy can initiate in the enteric nervous system and spread from the gut to the brain (*10-12*). Taken together, these data highlight gastrointestinal disruptions as notable prodromal symptoms of PD and raise the question of whether they may causally modify the risk for or manifestation of PD.

Mutations in the large multidomain protein leucine rich repeat kinase 2 (**LRRK2**) are associated with increased risk for both PD and IBD (*13*). The G2019S mutation in the kinase domain, which increases kinase activity, is the most common monogenic risk factor for PD, whereas the N2081D mutation, which is also in the kinase domain and increases kinase activity, is associated with IBD (*13*). Carriers of LRRK2 G2019S are at greater risk for developing Lewy body neuropathology, as well as motor deficits characteristic of PD (*14*). However, the penetrance of LRRK2 G2019S is incomplete and highly variable across study populations, suggesting that additional risk factors may interact with LRRK2 G2019S to cause PD (*15*). Whether the PD-associated G2019S mutation may influence susceptibility to intestinal inflammation and how these interactions may alter the onset and severity of neuropathological and behavioral symptoms of PD remains unclear.

In addition to intestinal inflammation, biological sex and age are factors of particular interest, given that PD is a progressive aging-related disorder that exhibits greater prevalence, earlier onset, and more severe motor impairment in males relative to females (*16*). Consistent with this, males with LRRK2 mutations display younger age of PD onset compared to males with idiopathic PD (*17*). However, this is contrasted by a separate study reporting younger age of onset for female LRRK2 mutation carriers relative to male carriers (*18*). Exactly how biological sex may influence interactions between intestinal inflammation, LRRK2 G2019S, and risk for PD is poorly understood. Herein, we dissect roles for sex as a biological variable that modifies gene-environment interactions between LRRK2 G2019S and prodromal intestinal inflammation.

To do so, we first modeled prodromal intestinal inflammation in pre-symptomatic male vs. female transgenic mice expressing human LRRK2 G2019S (**hLRRK2**^**G2019S**^ **Tg**, (*19*)) by subjecting 10-13 week old mice to 3 rounds of 2% dextran sulfate sodium (**DSS**) in water to induce intestinal damage as a common model of experimental colitis (*20*) (**Fig. 1A**). Compared to wildtype (**WT**) littermate controls, both male and female hLRRK2^G2019S^ Tg mice exhibited hypersensitivity to DSS-induced intestinal inflammation, as measured by more severe decreases in body weight relative to WT controls (**Fig. 1B**). While both male and female hLRRK2^G2019S^ Tg mice exhibited weight loss of a similar degree, male hLRRK2^G2019S^ Tg mice exhibited a delayed initial response relative to female hLRRK2^G2019S^ Tg mice (**Fig. 1B**), with more severe clinical score, which considers initial weight loss, stool consistency, and rectal bleeding ((*21*), **Fig. 1E**), and reduced survival (**Fig. 1F**). The genotype-dependent exacerbation of DSS response was most notable during the first round of DSS treatment (**Fig. 1B**), with diminished effects in the second round of DSS treatment (**Fig. 1C**), and no significant effect during the third round of DSS treatment (**Fig. 1D**), suggesting that LRRK2 G2019S may impact early immune responses to DSS-induced intestinal damage (*22*). Consistent with this, we observed a genotype-dependent increase in serum IL-12p40, IL-9, and IL-33 after 7 days of DSS treatment (**Fig. 1, G to I**), with no significant change in other conventional pro-inflammatory cytokines such as TNFa (**fig. S1A**). Colonic CD64^+^ MHCII^+^CD11b^+^ macrophages were also elevated in DSS-treated hLRRK2^G2019S^ Tg mice compared to the WT littermates (**Fig. 1J**), with no significant alterations in other colonic lamina proprial populations, such as Ly6C^+^ monocytes, CD103^+^ dendritic cells, T-helper (**Th**)1 cells, and Th17 cells (**fig. S1, B to E**). Overall, these results reveal that male and female hLRRK2^G2019S^ Tg mice exhibit increased sensitivity to intestinal damage during the acute phase of DSS exposure.

**Fig. 1:**
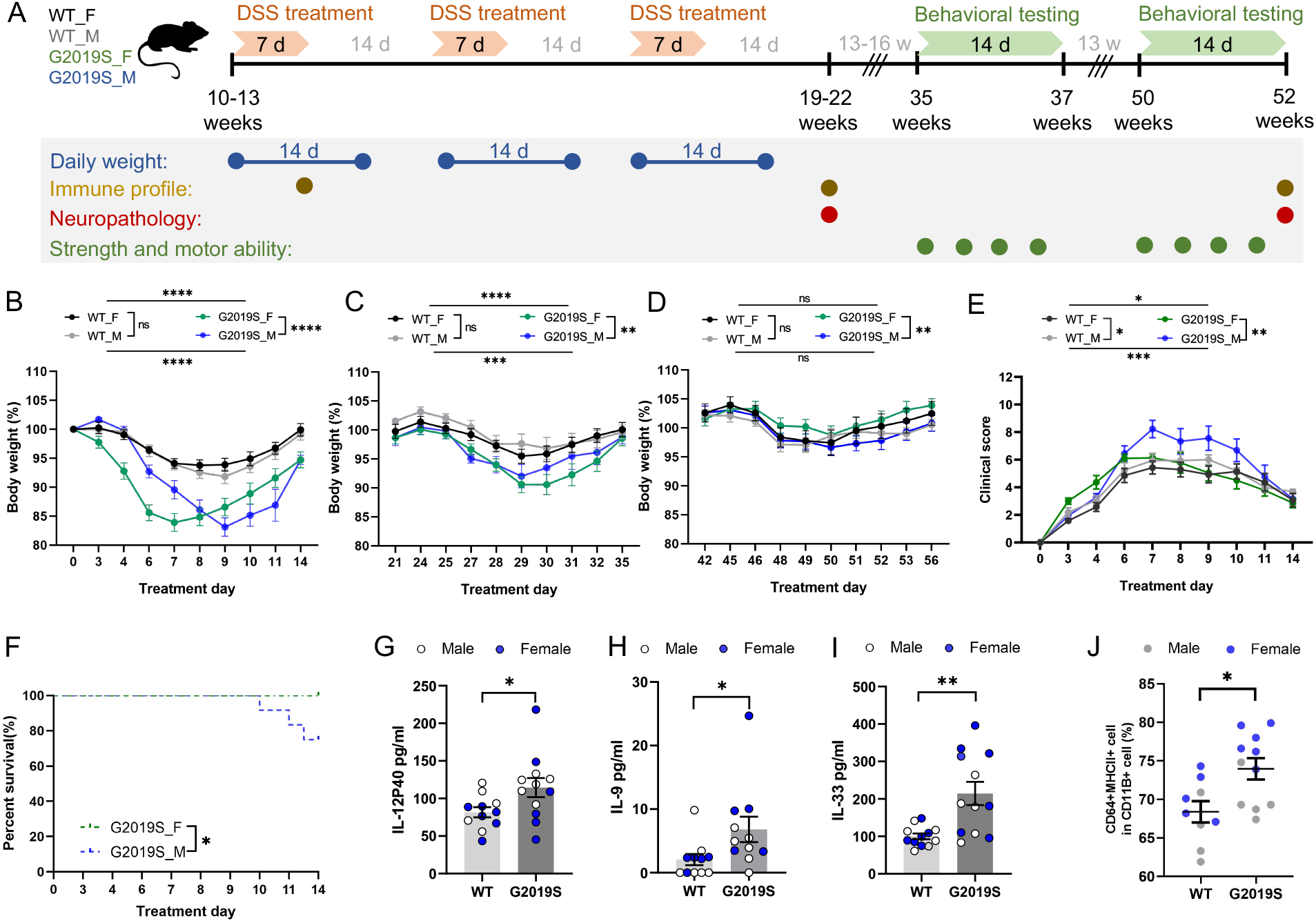
hLRRK2^G2019S^ Tg mice exhibit aggravated acute responses to DSS-induced intestinal inflammation. **(A)** Experimental timeline **(B-D)** Body weight normalized to starting weight (day 0) for male (M) and female (F) hLRRK2^G2019S^ mice (G2019S) and wildtype (WT) littermate controls over 3 rounds DSS treatment (two-way ANOVA with Sidak, n=12-18). **(E)** DSS clinical score over first round of treatment (two-way ANOVA with Sidak, n=7-9). **(F)** Survival curves (Survival analysis, n=12-18). **(G-I)** Concentrations of serum pro-inflammatory cytokines on day 7 of DSS treatment (two-tailed t-test, n=9-11). **(J)** Proportion of colonic lamina proprial macrophages on day 7 of DSS treatment. (two-tailed t-test, n=9-11). Data are presented as mean ± SEM. \**P*<0.05; \*\**P*<0.01; \*\*\**P*<0.001; \*\*\*\**P*<0.0001.

hLRRK2^G2019S^ Tg mice develop aging-dependent motor impairments that are detectable by behavioral testing beginning at 65 weeks of age (*23*). To determine the effect of prodromal intestinal inflammation on the onset and severity of LRRK2 G2019S-driven motor impairments, 10-13 week old male and female hLRRK2^G2019S^ Tg mice were subjected to 3 rounds of DSS treatment as model of chronic experimental colitis (*20*), and then evaluated in a battery of motor behavioral tests at ∼15 and ∼30 weeks after DSS treatment, reflecting 35 and 50 weeks of age (**Fig. 1A**). Compared to male WT littermate controls treated with DSS, male hLRRK2^G2019S^ Tg mice exhibited striking motor abnormalities in response to prodromal intestinal inflammation, including hyperactivity in the open field (**Fig. 2, A and C**) and reduced grip strength (**Fig. 2, B and C**), at both the 35- and 50-week old time points. This was not observed in DSS-treated female hLRRK2^G2019S^ Tg mice, which displayed only mild motor deficits in select metrics of the pole descent test and open field test, with results that were inconsistent between the two time points (**Fig. 2, A and C, fig. S2**). Notably, motor impairments were also largely absent in age-matched male hLRRK2^G2019S^ Tg mice that were treated with vehicle (water), instead of DSS (**Fig. 2D, fig. S3**), indicating that prodromal DSS-induced intestinal inflammation and the LRRK2 G2019S genotype interact to expedite and exacerbate motor abnormalities particularly in males.

**Fig. 2:**
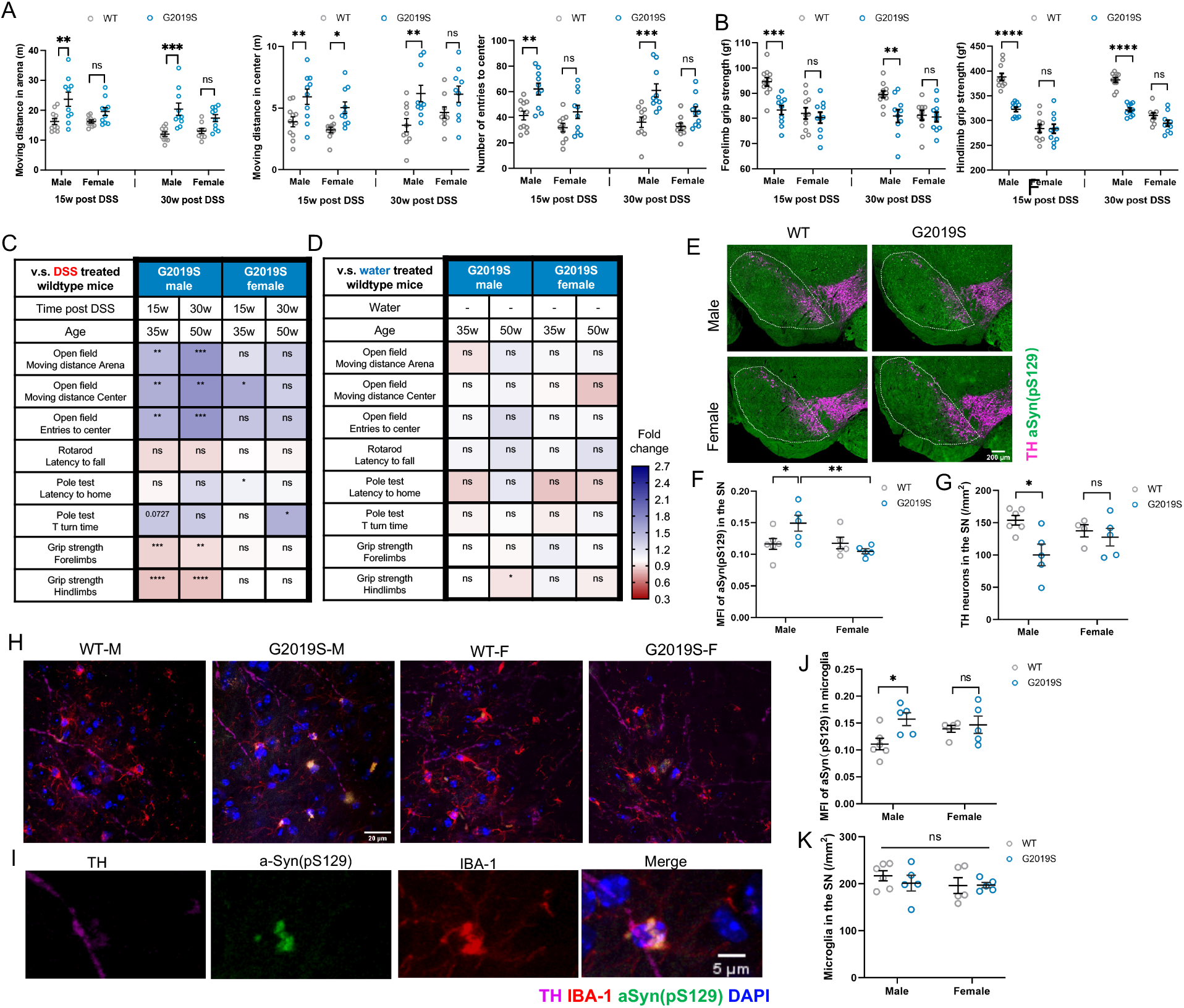
DSS-induced intestinal inflammation exacerbates behavioral and neuropathological features of PD in male hLRRK2^G2019S^ Tg mice. **(A)** Open field test: moving distance in the arena (left), center (middle) and number of entries to the center (right) after 10 minutes of exploration (two-way AVOVA with Sidak; n=9-10). **(B)** Grip strength for forelimbs (left) and hindlimbs (right) (two-way AVOVA with Sidak; n=9-10). **(C)** Summary of behavioral testing results for DSS-treated hLRRK2^G2019S^ Tg mice relative to DSS-treated WT littermate controls. **(D)** Summary of behavioral testing results for vehicle-treated (untreated) hLRRK2^G2019S^ Tg mice relative to WT littermate controls. **(E)** Representative images of TH+ neurons (magenta) and a-Syn (green) in the substantia nigra (SN) of hLRRK2^G2019S^ Tg mice and WT littermates at 32 weeks post DSS treatment (52 weeks of age). Scale bar = 200 um. **(F)** Mean fluorescence intensity (MFI) of a-Syn in the SN (two-way ANOVA with Sidak, n=5) **(G)** Number of TH+ neurons in the SN (two-way ANOVA with Sidak, n=5). (H, I) Representative images of IBA-1^+^ microglia (red) and a-Syn (green) in the substantia nigra (SN) of male and female hLRRK2^G2019S^ Tg mice and WT littermates at 32 weeks post DSS treatment (52 weeks of age). **(J)** MFI of a-Syn in IBA-1^+^ microglia in the SN (two-way ANOVA with Sidak, n=5). **(K)** Number of IBA-1^+^ microglia in the SN (two-way ANOVA with Sidak, n=5). Data are presented as mean ± SEM. \**P*<0.05; \*\**P*<0.01; \*\*\**P*<0.001; \*\*\*\**P*<0.0001.

To further assess effects of prodromal intestinal inflammation, LRRK2 G2019S, and biological sex on PD-related neuropathology, 10-13 week old male and female hLRRK2^G2019S^ Tg mice and WT littermates were subjected to DSS treatment and assessed 30 weeks later (at 50 weeks of age) for α-synuclein aggregation and dopaminergic neuron loss in the substantia nigra pars compacta (**SN**), two hallmark pathologies of PD (*24*). DSS-treated male hLRRK2^G2019S^ Tg mice exhibited elevated α-synuclein aggregation in the SN compared to DSS-treated male WT littermates (**Fig. 2, E and F**), whereas DSS-treated female hLRRK2^G2019S^ Tg mice exhibited no difference relative to their female WT controls. Consistent with the increases in α-synuclein burden, decreased numbers of tyrosine hydroxylase (**TH**^**+**^) neurons were detected in the SN of DSS-treated male hLRRK2^G2019S^ Tg mice compared to their treatment-matched WT controls, with no genotype-dependent differences in females (**Fig. 2G**). These neuropathological abnormalities were not seen in male hLRRK2^G2019S^ Tg mice treated with vehicle (water) instead of DSS (**fig. S4**), indicating that prodromal intestinal inflammation expedites the onset and severity of PD-related neuropathology in genetically predisposed males.

We observed that some microglia from DSS-treated hLRRK2^G2019S^ Tg mice contained α-synuclein deposits and were localized adjacent to TH^+^ axons (**Fig. 2, H and I**). Given that microglia are able to internalize neuron-derived α-synuclein (*25*) and accumulate α-synuclein intracellularly to promote dopaminergic neuron loss (*26*), we quantified α-synuclein loads localized within IBA-1^+^ microglia. Mean fluorescence intensity (**MFI**) of α-synuclein within SN microglia was elevated in DSS-treated male hLRRK2^G2019S^ Tg mice compared to DSS-treated male WT littermates (**Fig. 2J)**, with no differences in the absolute numbers of IBA-1^+^ microglia in the SN (**Fig. 2K**). Both male and female hLRRK2^G2019S^ Tg mice and WT littermates exhibited increased IBA-1 and CD68 intensity in the SN in response to DSS treatment (**fig. S5**), indicating that prodromal intestinal inflammation broadly stimulates microglial activation in both biological sexes, independently of genotype. Taken together, these findings reveal that prodromal intestinal inflammation potentiates adverse effects of LRRK2 G2019S in a sex-dependent manner, to promote α-synuclein aggregation in the SN, α-synuclein accumulation within SN microglia, loss of dopaminergic neurons, and impaired motor behavior, particularly in genetically predisposed males.

To further assess the specificity of this gene-environment interaction to the G2019S mutation in human LRRK2, we tested the effects of prodromal intestinal inflammation in two additional mouse lines—mice harboring a knock-in (**KI**) G2019S mutation in the endogenous mouse LRRK2 (**mLRRK2**^**G2019S**^ **KI**, (*27*)) and transgenic mice expressing human WT LRRK2 (**hLRRK2**^**WT**^, (*28*)). In contrast to the increased sensitivity of hLRRK2^G2019S^ Tg mice to acute DSS treatment (**Fig. 1**), mLRRK2^G2019S^ KI mice exhibited DSS-induced body weight loss that was mostly comparable to that seen in WT littermate controls (**fig. S6, A to C**). Despite no overt differences in responsiveness to prodromal intestinal inflammation, both male and female mLRRK2^G2019S^ KI mice treated with DSS went on to develop motor deficits in the open field, rotarod, pole descent, and grip strength tests at 15 and 30 weeks after DSS treatment, as compared to treated WT littermate controls (**fig. S6, D to H**). These motor impairments were largely absent in male and female mLRRK2^G2019S^ KI mice that were treated with vehicle (water) instead of DSS (**fig. S7**), indicating that prodromal intestinal inflammation expedites and/or exacerbates motor impairment induced by the G2019S mutation in mouse LRRK2.

Although overexpression of WT LRRK2 is used to mimic the elevated kinase activity induced by the G2019S mutation (*28*), hLRRK2^G2019S^ and hLRRK2^WT^ Tg mice demonstrated distinct responses to DSS-induced intestinal inflammation. While hLRRK2^G2019S^ mice displayed increased sensitivity to DSS treatment compared to treated WT controls (**Fig. 1**), hLRRK2^WT^ Tg mice exhibited modest reductions in body weight loss in response to acute DSS treatment, with males showing poor recovery relative to females and WT controls (**fig. S8, A to C**). Despite these differential acute responses to DSS treatment, both male and female hLRRK2^WT^ Tg mice developed motor impairments in response to prodromal intestinal inflammation that were detectable at 15 weeks and 30 weeks after DSS treatment (35 and 50 weeks of age), with male hLRRK2^WT^ Tg mice exhibiting more severe behavioral defects than females, relative to their respective sex- and treatment-matched WT littermate controls (**fig. S8, D to H**). Notably both mLRRK2^G2019S^ KI and hLRRK2^WT^ Tg developed genotype- and DSS-dependent motor impairments despite no increases in sensitivity to DSS treatment, indicating that the severity of response to DSS-induced intestinal inflammation does not correlate with the severity of motor impairment; rather, DSS treatment may serve as an environmental trigger of mechanisms that expedite or enhance pathogenesis in the context of genetic predisposition. Moreover, the bias for genetically predisposed males to develop more severe and earlier onset motor impairment in response to prodromal intestinal inflammation was seen only in mice expressing human LRRK2, and human LRRK2 G2019S especially, which may be due to transgenic overexpression of the proteins or species-specific differences in the activity of human vs. mouse LRRK2 (*29*). Altogether, findings from these experiments reveal that early DSS treatment reproducibly promotes the onset and severity of motor impairment across three different LRRK2-based mouse models for PD (**Supplemental Text**). These results strongly suggest that gut-brain interactions between prodromal intestinal inflammation and LRRK2 G2019S elevate risk for PD.

The heightened susceptibility of male hLRRK2^G2019S^ Tg mice to PD-related motor impairments and neuropathological abnormalities in response to DSS treatment (**Fig. 1, 2**) presents the opportunity to identify mechanisms underlying the ability of biological sex to modify risk for PD. To gain insight, we first asked whether there are sex differences in the expression of endogenous mouse LRRK2 or human LRRK2 G2019S in male vs. female hLRRK2^G2019S^ Tg mice and WT littermate controls, at baseline or 7 days after DSS treatment. In contrast to a prior report that LRRK2 expression is increased in inflamed tissues (*30*), we observed no change in the expression of endogenous mouse LRRK2 transcript in the colon and decreased expression of transgenic human LRRK2 transcript in the colon on day 7 of DSS treatment, with no sex differences at baseline or after DSS treatment (**fig. S9**). These findings suggest that the observed sex differences in response to prodromal intestinal inflammation are not due to differential expression of *LRRK2* between males and females.

To further assess the potential for gonadal hormones to mediate the observed sex differences in risk for PD-related symptoms (*31*), we performed gonadectomies in male hLRRK2^G2019S^ Tg mice 3 weeks before DSS treatment to evaluate the necessity of testicular hormones to exacerbate acute responses to experimental colitis and expedite the onset of motor impairment (**Fig. 3A**). Gonadectomy in male hLRRK2^G2019S^ Tg mice prevented the enhanced body weight loss seen during the acute phase of DSS treatment in male hLRRK2^G2019S^ Tg mice that were untreated (**Fig. 1B**) or subjected to a mock surgical procedure (**Mock**) (**fig. S10A**). Despite the normalized body weight response to DSS-induced intestinal inflammation, gonadectomized (**GDX**) and DSS-treated hLRRK2^G2019S^ Tg mice still developed motor impairments analogous to those seen in Mock DSS-treated hLRRK2^G2019S^ Tg controls (**Fig. 3, B to D; fig. S10, B and C**). Consistent with conclusions drawn from the experiments with mLRRK2^G2019S^ KI and hLRRK2^WT^ Tg mice, this again decouples the severity of acute DSS response from the severity and onset of motor impairment, highlighting a role for DSS-induced intestinal inflammation as an environmental trigger that potentiates genetic risk for PD symptoms in LRRK2 carriers (**Supplemental Text**). Furthermore, the results indicate that gonadal hormones are not required for the ability of prodromal intestinal inflammation to potentiate motor impairments in male LRRK2 G2019S carriers.

**Fig. 3:**
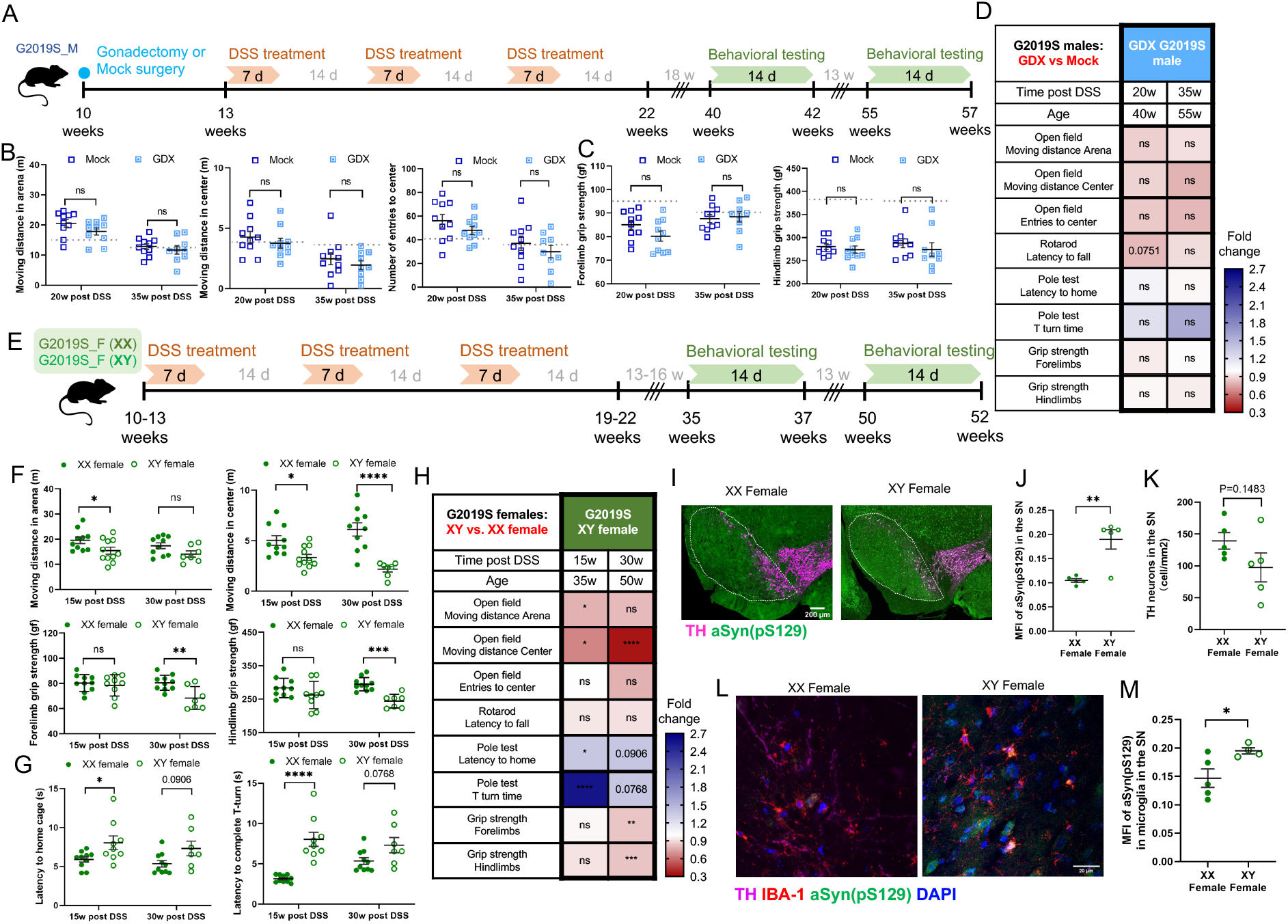
Male sex chromosomes, but not gonadal hormones, promote behavioral and neuropathological features of PD in hLRRK2^G2019S^ Tg mice exposed to early DSS-induced intestinal inflammation. **(A)** Experimental timeline. **(B)** Open field test: moving distance in the arena (left), center (middle) and number of entries to the center (right) after 10 minutes of exploration (two-way ANOVA with Sidak, n=9-10). **(C)** Grip strength for forelimbs (left) and hindlimbs (right) of gonadectomized (GDX) or mock surgery-exposed (Mock) male hLRRK2^G2019S^ Tg mice at 20 and 35 weeks after DSS treatment (two-way ANOVA with Sidak, n=9-10). Dotted lines indicate the mean value for WT littermates as a reference. **(D)** Summary of behavioral testing results for male GDX hLRRK2^G2019S^ Tg mice at 20 and 35 weeks post DSS treatment relative to male Mock hLRRK2^G2019S^ Tg controls (two-way ANOVA with Sidak, n=9-10). **(E)** Experimental timeline. **(F) Top:** Open field test: moving distance in the arena (left), center (middle), and number of entries to the center (right) after 10 minutes of exploration (two-way ANOVA with Sidak, n=7-9). Bottom: Grip strength for forelimbs (left) and hindlimbs (right) (two-way ANOVA with Sidak, n=7-9). **(G)** Pole descent test: latency to return to the home cage (left) and latency to complete T turn on the pole (right) for XX and XY female hLRRK2^G2019S^ Tg mice at 15 and 30 weeks post DSS treatment (two-way ANOVA with Sidak, n=7-9). **(H)** Summary of behavioral testing results for XY female hLRRK2^G2019S^ Tg mice at 15 and 30 weeks after DSS treatment relative to XX female hLRRK2^G2019S^ Tg controls (two-way ANOVA with Sidak, n=7-9). **(I)** Representative images of TH^+^ neurons (magenta) and a-Syn (green) in the substantia nigra (SN) of XX and XY female hLRRK2^G2019S^ Tg mice at 32 weeks post DSS treatment (52 weeks of age). Scale bar = 200 um. **(J)** Mean fluorescence intensity (MFI) of a-Syn in the SN (two-tailed t-test, n=5) **(K)** Number of TH+ neurons in the SN (two-tailed t-test, n=5). (**L**) Representative images of IBA-1^+^ microglia (red) and a-Syn (green) in the substantia nigra (SN) of XX and XY female hLRRK2^G2019S^ Tg mice at 32 weeks post DSS treatment (52 weeks of age). **(M)** MFI of a-Syn in IBA-1^+^ microglia in the SN (two-tailed t-test, n=5). Data are presented as mean ± SEM. \**P*<0.05; \*\**P*<0.01; \*\*\**P*<0.001; \*\*\*\**P*<0.0001.

To evaluate roles for sex chromosomes in causing greater male risk for PD-related behavioral and neuropathological symptoms in response to prodromal intestinal inflammation (*32*), we mated XX female hLRRK2^G2019S^ Tg mice with XY^-^*Sry* gonadal male mice in which the testis determining gene *Sry* was removed from the Y chromosome and inserted autosomally (*32, 33*). In so doing, we generated XY and XX gonadal female carriers of the human LRRK2 G2019S transgene. XY and XX gonadal female hLRRK2^G2019S^ Tg mice were subjected to 3 rounds of DSS treatment (**Fig. 3E**). Both XX and XY gonadal female hLRRK2^G2019S^ Tg mice exhibited severe body weight loss during the acute phase of DSS treatment (**fig. S11A**), which was comparable to the genotype-dependent enhancements in body weight loss seen in conventional female hLRRK2^G2019S^ Tg mice (**Fig. 1B**). Notably, compared to XX controls, XY gonadal female hLRRK2^G2019S^ Tg mice also exhibited delayed body weight loss (**fig. S11A**), similar to that seen in male hLRRK2^G2019S^ Tg mice treated with DSS (**Fig. 1B**). This indicates that conferral of XY sex chromosomes to gonadal female hLRRK2^G2019S^ Tg mice phenocopies the body weight responses to DSS treatment seen in male hLRRK2^G2019S^ Tg mice.

To further assess roles for sex chromosomes in regulating risk for motor impairment, DSS-treated XY and XX gonadal female hLRRK2^G2019S^ Tg mice were subjected to behavioral testing in a battery of motor tasks at 15 and 30 weeks after DSS treatment (35 and 50 weeks of age). DSS-treated XY gonadal female hLRRK2^G2019S^ Tg mice exhibited decreased moving distance in the open field, weaker grip strength, and longer latency to maneuver in the pole descent test, compared to treatment-matched XX hLRRK2^G2019S^ Tg littermate controls (**Fig. 3, D to H; fig. S11, B and C**). We further examined the PD pathology in the brains of XY female hLRRK2^G2019S^ Tg mice after behavioral testing (52 weeks of age). DSS-treated XY female hLRRK2^G2019S^ Tg mice exhibited significantly increased levels of α-synuclein aggregation in the SN relative to DSS-treated XX hLRRK2^G2019S^ Tg controls (**Fig. 3, I and J**). Increased burden of α-synuclein was also seen in SN microglia of DSS-treated XY female hLRRK2^G2019S^ Tg mice relative to treatment-matched XX controls (**Fig. 3, L and M**), which phenocopies the sex-dependent neuropathological phenotypes seen in male hLRRK2^G2019S^ Tg mice exposed to prodromal intestinal inflammation (**Fig. 2, E to I**). There were modest, but not statistically significant, decreases in TH^+^ neurons in DSS-treated XY female hLRRK2^G2019S^ Tg mice relative to XX controls (Fig. 3K). Overall, these results reveal that differences in effects of XY vs. XX sex chromosomes, not testicular hormones, likely cause sex differences in gene-environment interactions between prodromal intestinal inflammation and LRRK2 G2019S that increase risk for behavioral and neuropathological features of PD particularly in male hLRRK2^G2019S^ Tg mice (**Supplemental Text**).

To gain insight into the early gut-brain interactions that enable prodromal intestinal inflammation to promote LRRK2 G2019S-driven risk for motor impairment, α-synuclein aggregation, and dopaminergic neuron loss, we first evaluated DSS-induced alterations in the colon, serum, and brain after the 3^rd^ round of DSS treatment, before the onset of motor deficiencies (∼19 weeks of age, **Fig. 1A**). In light of evidence that intestinal seeding of α-synuclein can promote protein misfolding and spread to the brain (*10-12*), we quantified levels of α-synuclein in the colon of DSS-treated male and female hLRRK2^G2019S^ Tg mice. Compared to vehicle-treated hLRRK2^G2019S^ Tg controls, male DSS-treated hLRRK2^G2019S^ Tg mice exhibited increases in α-synuclein levels in colonic tissue (**Fig. 4A**) and in α-synuclein^+^ macrophages in the colonic lamina propria (**Fig. 4, B and C**). These alterations were not seen in female DSS-treated mice relative to their vehicle-treated controls, indicating a sex- and treatment-dependent enrichment in intestinal α-synuclein in hLRRK2^G2019S^ Tg mice. Consistent with the elevated α-synuclein loads detected in SN microglia of DSS-treated male hLRRK2^G2019S^ Tg mice at symptomatic ages (52 weeks, **Fig. 2, H to J**), increased α-synuclein levels were similarly detected in SN microglia of male, but not female, hLRRK2^G2019S^ Tg mice at a pre-symptomatic time point immediately after the third round of DSS treatment (∼19 weeks, **Fig. 4, D to F; fig. S12A**). There were no differences in levels of CD4^+^ T cells, CD8^+^ T cells or B cells in the brain parenchyma, or in various pro-inflammatory cytokines in the cerebrospinal fluid (CSF) of DSS-treated hLRRK2^G2019S^ Tg mice relative to vehicle-treated controls (**fig. S12, B to E**). Consistent with alterations in serum cytokines detected on day 7 of DSS treatment (∼11 weeks), serum IL-12p40 was similarly elevated in male and female hLRRK2^G2019S^ Tg mice after the 3 rounds of DSS treatment (∼19 weeks) (**fig. S12, F and H**), suggesting chronic elevation of this cytokine in response to DSS-induced intestinal inflammation. Serum IL-17A was also elevated at this time point (**fig. S12G**). However, there were no sex differences in the DSS-induced increases in select serum cytokines (**fig. S12H**). Taken together, these data indicate that DSS-induced intestinal inflammation leads to a sex-dependent accumulation of α-synuclein in colonic tissue, increase in α-synuclein^+^ macrophages of the colonic lamina propria, and elevation of α-synuclein loads within microglia of the SN, which all precede the sex-dependent onset of PD-related neuropathology and motor impairments in male hLRRK2^G2019S^ Tg mice. Overall, findings from this study indicate that early intestinal disruptions causally exacerbate pathology and motor dysfunction conferred by various genetic and environmental risk factors for PD (**Supplemental Text**).

**Fig. 4:**
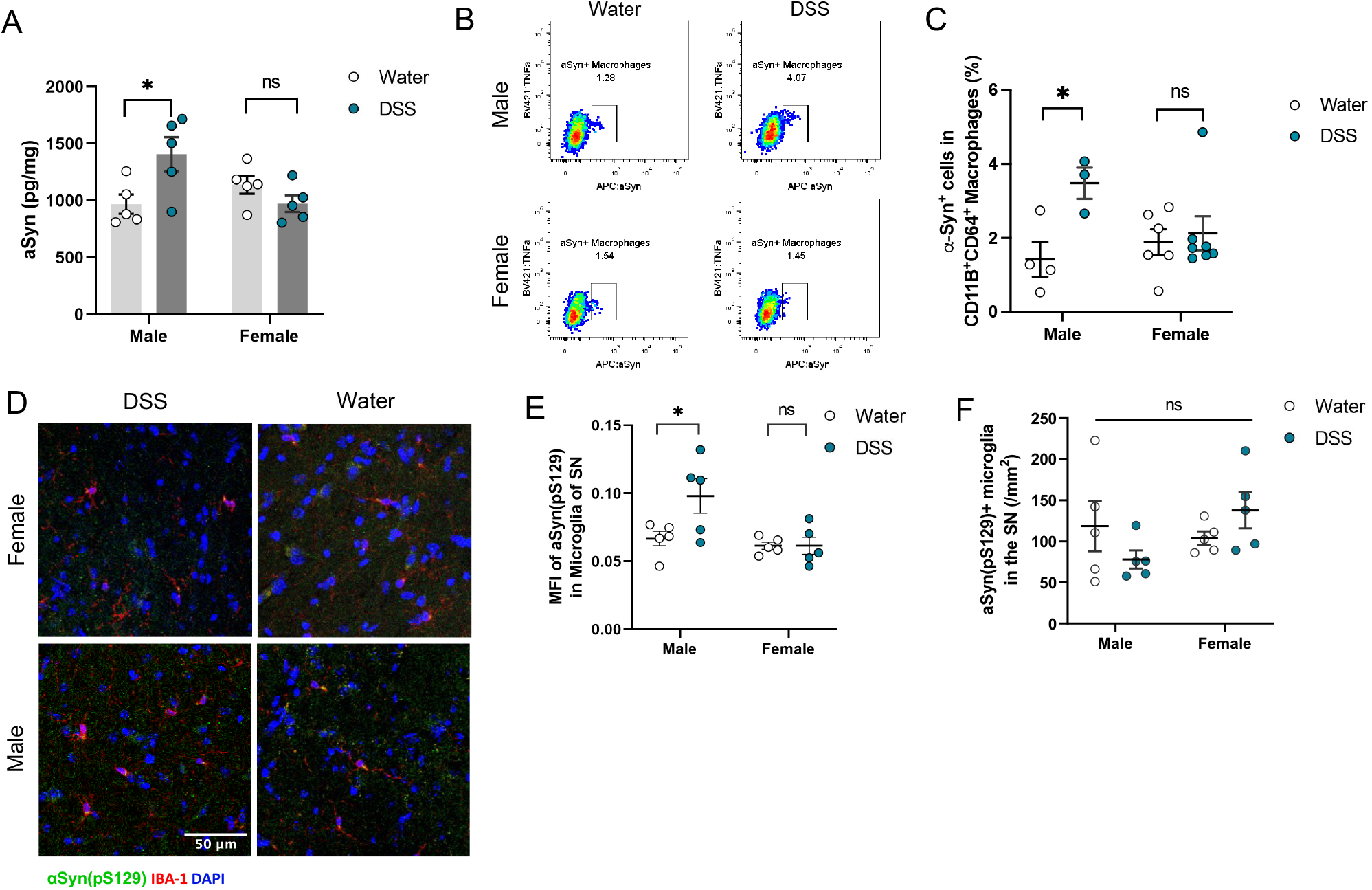
DSS-induced intestinal inflammation increases colonic a-Syn and a-Syn+ macrophages in pre-symptomatic male hLRRK2^G2019S^ Tg mice to promote the development of behavioral and neuropathological features of PD. **(A)** Levels of a-Syn detected in colons of hLRRK2^G2019S^ Tg mice at 1 week after the last round of DSS treatment (20-23 weeks of age) (two-way ANOVA with Sidak, n=5). **(B)** Representative flow cytometry plots for a-Syn^+^ macrophages from the colonic lamina propria of hLRRK2^G2019S^ Tg mice at 1 week after the last round treatment with DSS or standard water. **(C)** Quantitation of a-Syn^+^ macrophages from the colonic lamina propria of hLRRK2^G2019S^ Tg mice at 1 week after the last round of treatment with DSS or standard water (two-way ANOVA with Sidak, n=3-6). **(D)** Representative images of IBA-1^+^ microglia (red) and a-Syn (green) in SN the substantia nigra (SN) of male and female hLRRK2^G2019S^ Tg mice at 1 week after the last round of treatment with DSS or standard water. **(E)** Mean fluorescence intensity (MFI) of a-Syn in IBA-1^+^ microglia in the SN (two-way ANOVA with Sidak, n=5). **(F)** Number of a-Syn^+^ IBA-1^+^ microglia in the SN (two-way ANOVA with Sidak, n=5). Data are presented as mean ± SEM, \**P*<0.05

## Acknowledgments

We thank Dr. Malú Tansey and all members of the Hsiao lab for their helpful feedback on the study.

## Funding

Chan Zuckerberg Initiative Ben Barres Career Acceleration Award (EYH)

New York Stem Cell Foundation Robertson Neuroscience Investigator Award (EYH)

NIH HL131182 (APA)

## Author contributions

Conceptualization: PF, EYH

Methodology: PF, HE, LWY, GA, KL, YD, JL, HH

Funding acquisition: EYH

Resources: APA, EYH

Writing – original draft: PF, EYH

Writing – review & editing: PF, HE, LWY, GA, KL, YD, JL, HH, APA, EYH

## Data and materials availability

Data generated or analyzed during this study are included in this published article and its supplementary information files.

## Materials and Methods

### Mice

hLRRK2^G2019S^ Tg (JAX #018785), hLRRK2^WT^ Tg (JAX #012445, (*28*)), and mLRRK2^G2019S^ KI (JAX #030961, (*27*)) mice were purchased from Jackson Laboratories and bred in at the UCLA Center for Health Sciences barrier facility. XY^−^(*Sry*^+^) male mice, fathers of the “Four Core Genotypes” model (JAX #010905, (*34*)) were bred at UCLA. Animals were maintained on a 12-h light-dark cycle in a temperature-controlled environment, with autoclaved bedding. Sterile water and autoclaved standard chow (Lab Diet 5010) were provided *ad libitum*. For all experiments, wildtype littermates were used as controls. All experiments were performed in accordance with the NIH Guide for the Care and Use of Laboratory Animals using protocols approved by the Institutional Animal Care and Use Committee at UCLA.

### Gonadectomy

All surgical procedures were performed under sterile conditions. Prior to anesthesia, the mouse was injected with 0.1 ml carprofen and sterile lactated ringer’s (1:99). The mouse was then anesthetized with isoflurane, shaved at the site of incision (scrotum), administered purolube petroleum jelly on the eyes to prevent dehydration, and placed on a warm pad heated by circulating water bath. The scrotum was washed three times alternately with betadyne and 70% ethanol, and then a small cut was made in the scrotum and cremaster muscle. The testis was gently extruded through the incision site, clamped with a hemostat and ligated with an absorbable suture, then cut off. The hemostat was removed, the cremaster was then sutured, and the scrotum skin was shut with wound clips. The procedure was then repeated on the other side. The incision sites were treated with betadyne and topical antibiotic (Taro: polymyxin B sulfate, bacitracin zinc, and neomycin sulfate in petrolatum). The wound clips were removed 7-10 days later under light isoflurane anesthesia.

### DSS treatment

10-13-week old mice were given 2% DSS in autoclaved water for 1 week, followed by 2 weeks recovery, which was repeated an additional 2 times for a total of 3 rounds of DSS treatment to model prodromal ulcerative colitis (*20*). Body weight, stool consistency, and rectal bleeding were assessed daily during the 1 week of DSS treatment and 1 week following each DSS treatment. Clinical scores were calculated according to methods described in (*21*).

### Cytokine measurements

For CSF collection, mice were injected intraperitoneally with ketamine (100 mg/kg) and xylazine (10 mg/kg), and the cisterna magna was punctured according to methods detailed in (*35*). For serum collection, whole blood was collected from the aorta abdominalis, left at room temperature for 15 minutes to coagulate, and centrifuged at 2000g for 15 minutes. Serum and CSF samples were assessed for the cytokines IL-22, IFNg, IL-1b, IL-18, IL-23, IL-6, IL-4, IL-10, TGFb, TNFa, IL-33, IL-17A, IL-12p40, and IL-13 using a custom LEGENDplex kit according to manufacturer’s instructions (Biolegend). Flow cytometry was performed using a BD Celesta flow cytometer in UCLA’s Broad Stem Cell Research Center Flow Cytometry Core facility.

### Cellular immunoprofiling

Colon lamina propria leukocytes were isolated following procedures adapted from (*36*). Briefly, colons were aseptically dissected from mice anesthetized with isoflurane, rinsed twice in ice-cold PBS, and incubated while shaking for 20 min at 37°C in HBSS buffer containing 5mM EDTA and 10mM HEPES. Following 20 s vortex, supernatant was discarded and remaining tissues were incubated while shaking for 20 min at 37°C in pre-warmed RPMI buffer containing 4% FCS, 0.5 U/ml dispase, 0.5 mg/ml collagenase D, 0.25 mg/ml DNaseI. Following 20 s vortex, supernatant was discarded and tissues were minced in pre-warmed buffer and digested while shaking for an additional 45 min at 37°C. Following 20 s vortex, digested suspension was passed through a 70 um cell strainer, washed with ice cold RPMI, and centrifuged at 2000 rpm for 10 min at 4°C. Pelleted cells were resuspended in complete RPMI, stained with LIVE/DEAD Fixable Aqua Stain (Thermofisher) for 20 min following the manufacturer’s instructions, and then stained for subsets of markers at 5 *μ*g/ml: Ly6G-BV421(Biolegend, 127628), Ly6C-BV605 (Biolegend, 128036), CD4-BV650 (Biolegend, 100469), NK1.1-BV711 (Biolegend, 108745), B220-FITC (Biolgend, 103205), CD8a-PerCP-Cy5.5(Tonbo bioscience, 65-0081-U025), CD45-PE-Cy7 (Tonbo bioscience, 60-0451-U100), CD11c-APC-eFlour780 (Thermofisher, 47-0114-82), CD3-AF700 (Biolegend, 100215), CD11b-PE-eFlour610 (Thermofisher, 61-0112-80), MHCII-FITC (Thermofisher, 11-5321-82), CD64-PE-Cy7 (Biolegend, 139313). For intracellular staining, cells were fixed and permeabilized using the Fixation/Permeabilization kit (BD Biosciences) following the manufacturer’s instructions, and then stained for markers at 5 *μ*g/ml for IL-17A-FITC (Thermofisher, 11-7177-81) and IFNg-PE (Thermofisher, 12-7311-81), 10 *μ*g/ml for TNFa-BV421 (Biolegend, 506327), IL-1b-PE (Thermofisher, 12-7114-82), α-synuclein-APC (Novus Biologicals, NBP1-05194APC). Stained cells were resuspended in PBS, filtered through a 0.45 um strainer, and assessed via a BD Fortessa flow cytometer.

Brain leukocytes were isolated following procedures adapted from (*37*). Briefly, anesthetized mice were transcardially perfused with PBS, intact brains were dissected, stored in Hibernate-A medium (Thermofisher), and then transferred into a dounce homogenizer containing HBSS supplemented with 10% FBS, glutamine, and pen/strep. Tissues were homogenized by passing with slide plunger 6-8 times, triturated using a 5 ml serological pipet 5-6 times, and then passed through a 70 um filter with an additional wash with buffer. Suspension was centrifuged for 5 min at 500 g, and supernatant was discarded. Cells were resuspended in 1 ml 30% percoll, inverted 4-5 times, and then centrifuged for 7 min at 500 g. Myelin layer was carefully decanted, pellet was resuspended in ice-cold PBS, and centrifuged for 5 min at 500 g. Finally, cells were resuspended in PBS and stained for markers described above.

### Behavioral testing

Behavioral testing for motor ability was conducted at the UCLA Behavioral Testing Core. On each testing day, mice were habituated to the testing room for at least 30 minutes before initiating. Behavioral equipments were cleaned with 70% ethanol and Accel disinfectant before and after each session. Each behavioral test was conducted with at least 3 days of recovery between different tests.

#### Grip strength

Mice were lowered over the grip strength meter (Chatillion DFE-010, Ametek), allowing only the forepaws or hindpaws to attach to the grid with torso remaining horizontal before recording the maximal grip strength value of the mouse that is displayed on the screen. Each mouse was tested 5 replicate times for forelimb and for hindlimb, and the average value for each was recorded.

#### Open field test

Mice were placed in the center of a 30 cm x 30 cm arena for 10 min, during which an overhead Basler Gig3 camera and TopScan 3.0 (Clever Systems Inc.) software was used to measure distance traveled, and the number of entries and duration of time spent in the central 15 cm square area.

#### Pole test

Mice were placed head down at the top of a 50 cm vertical pole with a diameter of 1 cm, mounted on a base stand that was placed in the home cage. Time to reach the ground of the home cage was scored manually as latency to home cage. Separately, mice were placed head up at the top of the pole and the time to complete a 180° turn to reorient head down was scored manually as time to complete T-turn. Each test was assessed 5 replicate times, and the average value for each was recorded.

#### Rotarod test

Mice were trained on the rotarod instrument (Rotamex 5, Columbus Instruments) for 5 min twice consecutively, starting at 0.5 rpm and ending with 10 rpm, with accelerations of 1 rpm every 20 s. On the following day, mice were tested for 5 min, starting at 1 rpm and ending at 30 rpm, with accelerations of 1 rpm every 10 s. Each mouse was tested 3 replicate times, and the average value was recorded.

### Neuropathological assessments

#### Immunofluorescence staining

Mice were anesthetized with isoflurane, and then transcardially perfused with PBS and 4% paraformaldehyde (PFA). Brains were dissected and post-fixed in 4% PFA for 24 hours before embedding in optimal cutting temperature compound (OCT) and frozen in -80°C. Brains were cryosectioned at 25 *μ*m thickness targeting the SN and striatum and collected in PBS. Brain sections were then immunolabeled using standard free-floating technique in 5% donkey serum by incubating with antibodies for tyrosine hydroxylase (TH), ionized calcium binding adapter molecule 1 (IBA-1), CD68, and α-synuclein for later visualization of dopaminergic neurons, microglia, lysosomes, and α-synuclein aggregations, respectively. Samples were then incubated with suitable fluorophore-conjugated secondary antibodies in 5% donkey serum. Nuclei were stained with DAPI using slide mounting medium. Primary antibodies used: TH: AvesLabs, THY, Chicken anti-mouse/hu/ra, 1:500. IBA-1: Fujifilm Wako Chemicals, 019-1974, Rabbit anti-mouse, 1:1000. IBA-1: Fujifilm Wako Chemicals, 011-27991, Goat anti-mouse, 1:1000. CD68 Bio-Rad, MCA1957T, Rat anti-mouse, 1:400. α-synuclein pSer129: Novus Biologicals, NBP2-61121, Rabbit anti-mouse, 1:500. Secondary antibodies used: Alexa Fluor 647 AffiniPure Donkey Anti-Chicken (Jackson ImmunoResearch, 703-605-155) 1:1000. Alexa Fluor 488 AffiniPure Donkey Anti-Rabbit (711-545-152) 1:1000. Alexa Fluor 594 AffiniPure Donkey Anti-Rat (Jackson ImmunoResearch, 712-585-153) 1:1000. Alexa Fluor® 647 AffiniPure Donkey Anti-Goat (Jackson ImmunoResearch, 705-605-003) 1:1000. DAPI: ThermoFisher, P36962, ProLong(tm) Diamond Antifade Mountant with DAPI.

#### Confocal Imaging and Image Analysis

Confocal microscopy was utilized to acquire images of substantia nigra (SN) of mouse brains using Zeiss confocal (LSM 700, 20x magnification) microscope and Zen Black software. Only brains with well-defined TH-positive neurons in SN (*38-40*), as well as uniform SN, were imaged so each animal had 3-5 imaged sections for counting and analysis. Using Fiji imaging software, SN area was outlined, and subsequent image analysis was performed in the defined SN area. Average area of SN was 0.94 mm^2^ for 52-week-old mice. TH-positive cells were manually counted by a researcher blinded to the experimental group, ensuring that DAPI signal was localized to the neuronal cell body. Neuron counts were then normalized by SN area to assess neuron density. IBA-1 positive cells and CD68-positive or α-synuclein^+^ cells that were overlapping with IBA-1 positive cells were manually counted by a researcher blinded to experimental groups. Only cells with clear IBA-1 signal, microglia morphology, and DAPI overlay were counted. Positive CD68 signal in the SN was only quantified when within the IBA-1 signal as puncta, signifying lysosomal formation within the microglia. Similarly, positive α-synuclein signal in the microglia was only quantified when puncta were present within the microglia. All manual quantifications of IBA-1 positive-cells, positive-CD68 and α-synuclein puncta within IBA-1 were normalized by SN area. Mean fluorescence intensity (MFI) of IBA-1 and CD68-positive signals in the SN area was calculated to assess extent of microglia activation and lysosomal formation. Positive IBA-1 fluorescent signals in the SN were selected and quantified, and the background fluorescence of the image subtracted to target only fluorescent signals of IBA-1. Background fluorescence was subtracted to target only fluorescent signals of CD68. Quantifications were normalized by tissue area to account for varying SN size. Similarly, MFI of α-synuclein within the SN and α-synuclein within the IBA-1 signal was calculated to assess the extent of α-synuclein presence in the entire SN area or when contained in microglia, respectively.

### *LRRK2* qPCR

Colon and striatal tissue was lysed in RLE buffer and RNA was extracted using the RNeasy Kit (Qiagen). cDNA was synthesized by qScript cDNA SuperMix kit (Quantabio). Each reaction was set up with 1 *μ*L of DNA sample, 5 *μ*L of SYBR green master mix (ThermoFisher Scientific). forward (*human lrrk2*: 5’-TGATTCTCGTTGGCACACAT-3’, *mouse lrrk2*:5’-GCACATGCTCTGTCCACTCT-3’) and reverse (*human lrrk2*:5’-GCCAAAGCATCAGATTCCTC-3’, *mouse lrrk2*: 5’-CATGGGCATGCTTCTGCATC-3’) primers (Integrated DNA Technologies) at a final concentration of 500 nM each, and ultrapure water (ThermoFisher Scientific) to the final volume of 20 *μ*L. Quantitative PCR with reverse transcription (qPCR) was performed on a QuantStudio 5 thermocycler (ThermoFisher Scientific) with the following conditions: initial denaturation at 50° for 2 min and 95°C for 10 min followed by 40 cycles each consisting of denaturation at 95°C for 15 sec, annealing at 60°C for 60 sec and maintain at 4°C at the last step.

### Intestinal α-synuclein ELISA

Colon lysates were prepared as described for Western blots above. Total protein levels were measured by BCA assay (Pierce) and a 1:00 dilution was used for quantifying α-synuclein using the mouse α-synuclein ELISA kit (Abcam) following the manufacturer’s instructions.

### Statistical Analysis

Statistical analysis was performed using GraphPad Prism. Significance was determined by two-way ANOVA followed by Sidak’s post hoc test for multiple group comparisons. Comparisons included analyses of effects for genotype and treatment, and genotype and sex. All data are represented as mean ± s.e.m. (standard error of the mean). \**P* < 0.05 was considered statistically significant. \*\**P* < 0.01. \*\*\**P* < 0.001. Notable non-significant differences are indicated in the figures by “n.s.”.

## Supplemental Text

Findings from this study reveal that prodromal intestinal inflammation promotes the manifestation of PD-related motor impairments across three different mouse models of LRRK2 genetic risk – endogenous mouse LRRK G2019S, transgenic human LRRK2 G2019S and transgenic human WT LRRK2 overexpression. In addition, a recent study of a different human LRRK2 G2019S transgenic mouse on the FVB background similarly reported that monthly DSS treatment for 5 months reduced exploration in the open field and promoted PD-related neuropathology (*41*). In transgenic mice expressing the PD-associated A53T mutation in human a-synuclein, 3 months of mild continuous DSS treatment promoted a-syn aggregation in both enteric neurons and the SN, increased the loss of dopaminergic neurons in the SN, and expedited the onset of motor behavioral abnormalities (*42*). In Pink1^-/-^ mice, intestinal infection with *Citrobacter rodentium* resulted in degeneration of dopaminergic axons in the striatum and motor dysfunction (*43*). Moreover, oral rotenone treatment induced intestinal inflammation, motor dysfunction, and neurodegeneration in mice, which was prevented by deletion of TLR4 (*44*). Overall, our findings contribute to increasing evidence that early intestinal disruptions can exacerbate pathology and motor dysfunction conferred by various genetic and environmental risk factors for PD.

A major point of distinction in the experimental paradigm used in this study is the relatively early exposure to DSS-induced intestinal injury (at ∼10 weeks of age), followed by a relatively long recovery period of >4 months, which contrasts previous studies that employ continuous DSS treatment and physiological evaluations immediately following treatment (*41, 42*). This design is intended to model prodromal intestinal inflammation that can precede motor impairments in PD by up to 20 years (*3*). Notably, we observe no evidence of persistent inflammation, as indicated by full recovery of clinical score and minimal change in serum cytokines. This is consistent with a previous report of transient effects of DSS treatment on neuroinflammation, where transcriptional responses in the brain were seen at 7, but not 14, days after DSS treatment (*45*). Moreover, our results from DSS treatment across three different LRRK2 mouse models, as well as the gonadectomy and sex chromosome complement conditions, reveal that the severity of acute inflammatory response to DSS-induced intestinal injury does not correlate with the severity of motor impairments and neuropathological abnormalities that develop later in life. Rather, the findings indicate that intestinal inflammation serves as an early environmental trigger that exacerbates genetic risk posed by LRRK2 G2019S.

While the ability of prodromal intestinal inflammation to expedite and exacerbate endophenotypes of PD is shared across the three LRRK2 models we studied, we observed a sex bias toward males only in mice that expressed the human variant of LRRK2. The differences may be due to the overexpression human LRRK2 protein by 2.5-3.5X in the mouse transgenics as compared to the mouse knock-in strain (*46*). There are also species-specific differences in the expression of LRRK2 that could contribute—while both mouse endogenous *LRRK2* and human *LRRK2* transgene are expressed in the SN, the human *LRRK2* transgene also causes LRRK2 to be expressed in other neuronal subtypes, that more closely resembles descriptions of LRRK2 expression in humans and non-human primates (*47*). Nonetheless, the robustness of the sex difference we observed enabled us to probe the biological basis of the sex-dependent gene-environment interaction between prodromal intestinal inflammation and human LRRK2 G2019S in expediting and exacerbating risk for PD-related symptoms particularly in males.

We found that the sex chromosome complement (XY vs. XX), rather than testicular hormones, mediate the ability of prodromal intestinal inflammation to predispose male LRRK2 carriers to earlier onset and worsened severity of PD-related symptoms relative to females. This aligns with the male bias in the incidence of PD (*19*) and the earlier onset of motor dysfunction in male PD patients compared to females (*48, 49*). While little is known regarding roles for sex chromosomes in risk for PD, the *SRY* gene on the Y chromosome has been associated with neuroinflammation, dopaminergic neuron loss, and mitochondrial degradation (*50*). Inhibition of *SRY* expression in the substantial nigra attenuated the motor impairments and dopaminergic neuronal loss in a 6-hydroxydopamine (6-OHDA)-induced rat model of PD (*50*). Any potential effects of *Sry* on PD mechanisms cannot, however, explain the XY vs. XX differences found here in *Sry*-negative gonadal female mice. Moreover, these findings revealing an important role for sex chromosomes in promoting risk for PD contrasts other animal studies reporting neuroprotective effects of estrogen in neurotoxin-based models of PD (*51-53*). Additional studies are needed to determine whether the sex chromosomal effect is caused by the number of X chromosomes (including X dose, X imprint, and indirect effects of X inactivation), or the presence of Y the chromosome, and to further identify specific X or Y candidate genes that are responsible (*34, 54*). The X-lined gene *Kdm6a*, which escapes X-inactivation, may be of interest, given its link to sex-biased risk for autoimmune diseases and Alzheimer’s disease (*55, 56*). Results from our study warrant further evaluation of sex differences in the prevalence of PD and co-morbid prodromal intestinal dysfunction, particularly for LRRK2 carriers, as well as future experiments that identify specific sex-linked genetic factors that contribute to PD risk.

Consistent with the adverse outcomes particularly in predisposed males, we observed that prodromal intestinal inflammation led to increased levels of colonic α-synuclein and α-synuclein^+^ colonic macrophages that preceded evidence of neuroinflammation, neuropathology, and motor impairment in male LRRK2 G2019S carriers. A recent study reported that microglia engulf neuronally-derived α-synuclein and transfer the protein across microglial cells to promote effective degradation and clearance of pathogenic α-synuclein (*57*). Notably, the ability to transfer α-synuclein was impaired in LRRK2 G2019S microglia, resulting in α-synuclein accumulation and poor survival. Given that microglia have functional features in common with peripheral macrophages (*58, 59*), additional experiments are needed to determine if the increases in colonic α-synuclein^+^ macrophages particularly in male LRRK2 G2019S carriers exposed to prodromal intestinal inflammation reflect a cellular accumulation of α-synuclein due to poor clearance. α-Synuclein misfolding and aggregation is a pathological hallmark of PD that has the capacity to propagate from the gut to the brain (*10*). The preponderance of studies on gut-to-brain transmission of PD pathology have focused on trans-neuronal spreading of pathogenic α-synuclein through vagal gut-brain circuits that project to the SN (*12, 60, 61*). Further investigation is warranted to assess the potential for gut macrophages to contribute to this process, by the transfer α-synuclein from gut macrophages to gut-innervating vagal neurons, and/or the potential for α-synuclein^+^ gut macrophages to directly infiltrate the brain in response to intestinal damage particularly in male LRRK2 G2019S carriers.

---

**Fig. S1:**
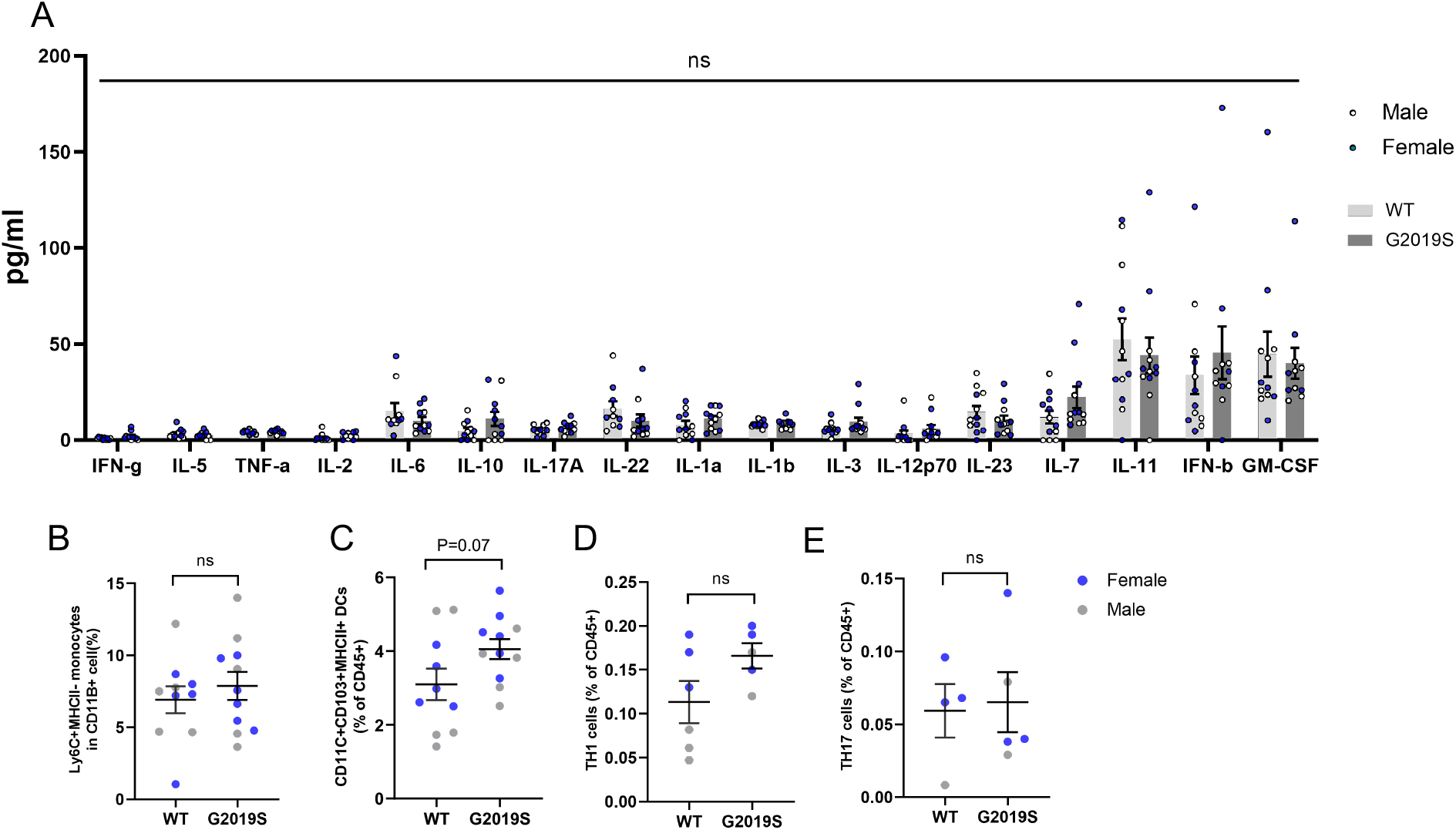
Serum cytokines and colonic lamina proprial immune cell subsets in response to DSS-induced intestinal inflammation in hLRRK2^G2019S^ Tg mice. (**A**) Serum cytokines in hLRRK2G2019S Tg mice (G2019S) relative to wildtype (WT) littermate controls on day 7 of DSS treatment (two-tailed t-test, n=12). (**B**) Monocytes, (**C**) dendritic cells, (**D**) TH1 cells and (**E**) TH17 cells from the colonic lamina propria of G2019S vs. WT mice on day 7 of DSS treatment (two-tailed t-test, n=4-10). Data are represented as mean ±SEM; ns= not statistically significant.

**Fig. S2:**
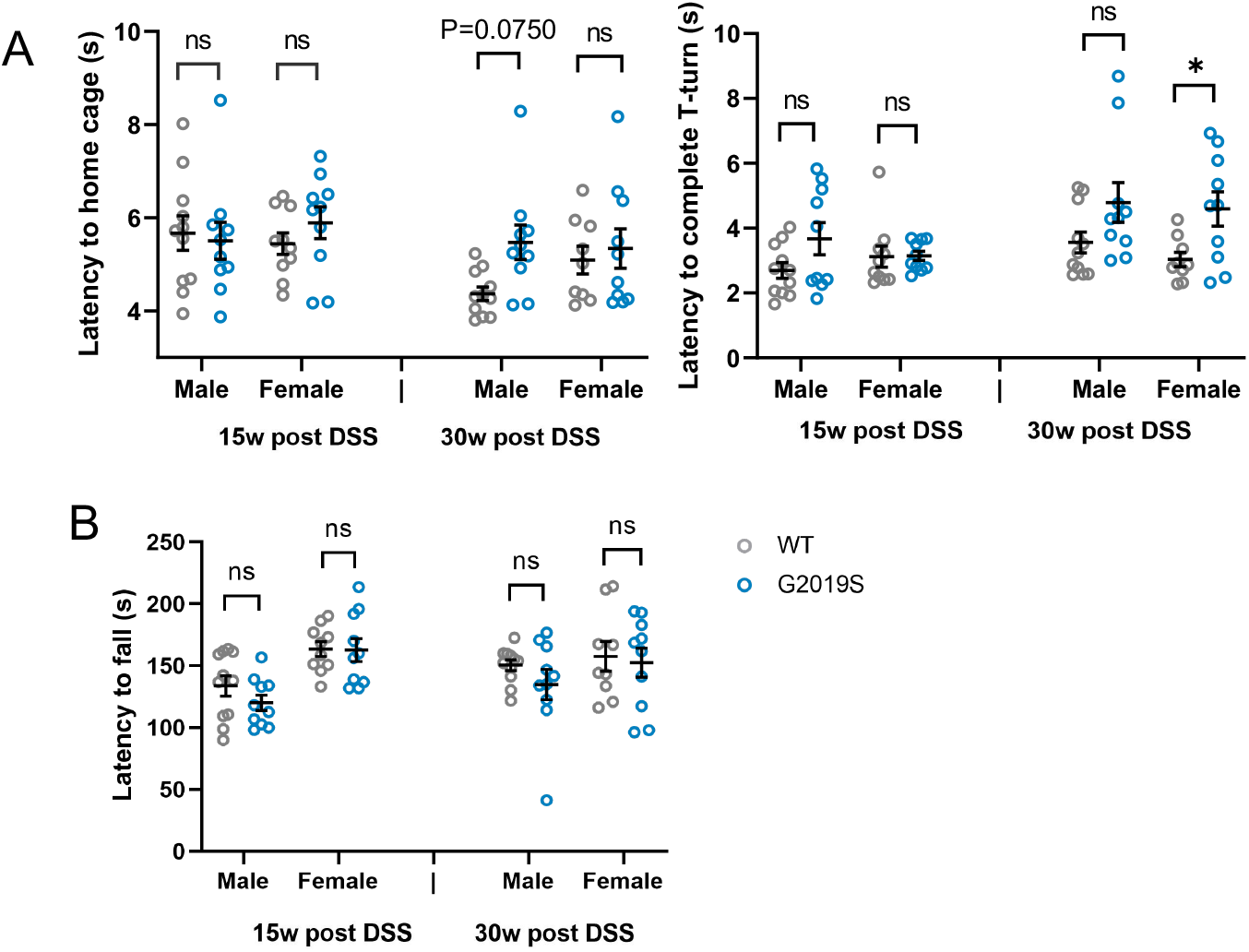
Behavioral performance in additional motor tasks for hLRRK2^G2019S^ Tg mice after prodromal DSS-induced intestinal inflammation. (**A**) Pole descent test: latency to return to the home cage (left) and latency to complete T turn on the pole (right). (**B**) Rotarod test for hLRRK2^G2019S^ Tg mice (G2019S) compared to wildtype (WT) littermate controls at 15 and 30 weeks post DSS treatment (Two-way ANOVA with Sidak, n=9-10). Data are presented as mean ±SEM, \**P*<0.05.

**Fig. S3:**
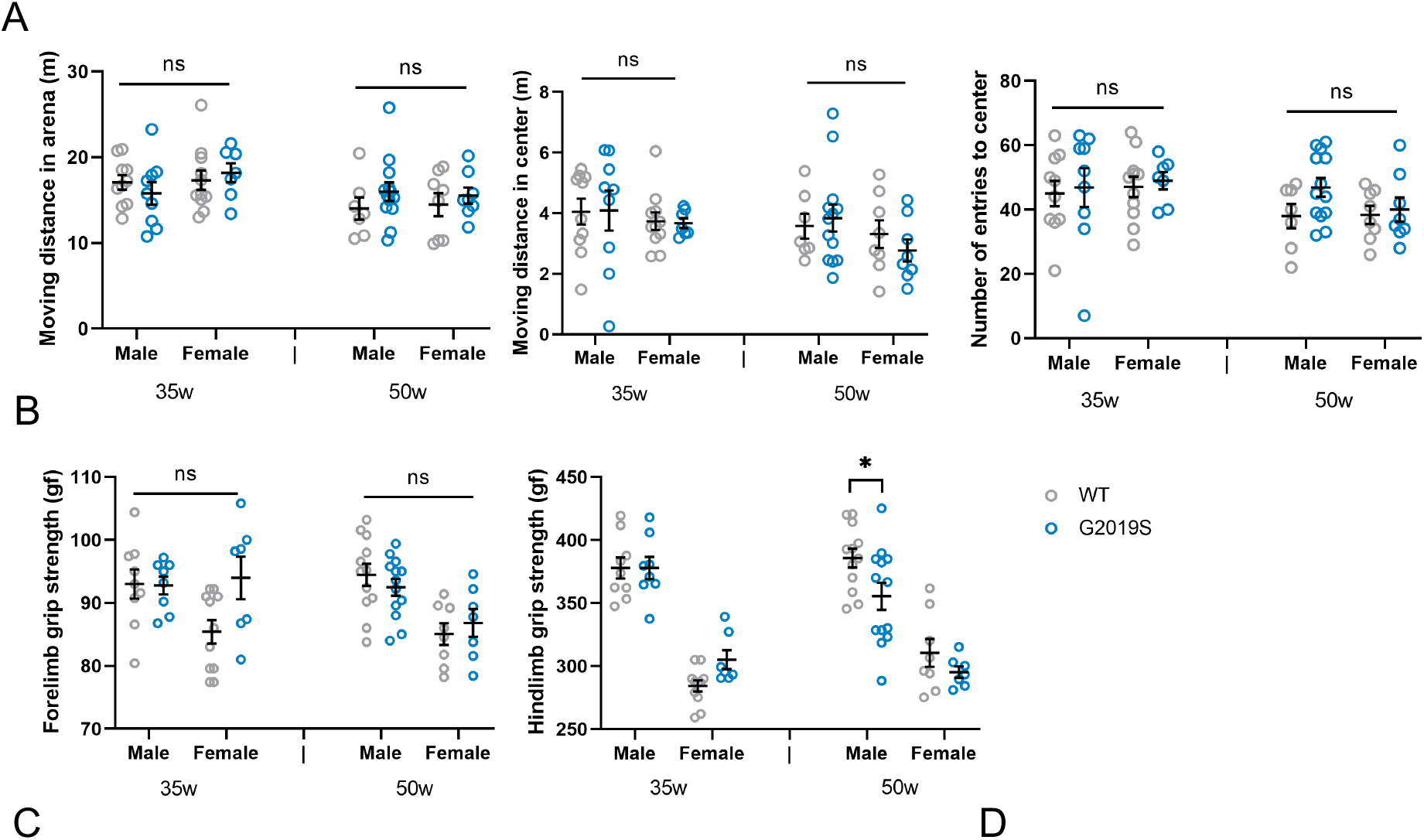
Vehicle-treated (water; untreated) hLRRK2^G2019S^ Tg mice and WT littermate controls in behavioral tests for motor ability. (**A**) Open field test: moving distance in the arena (left), center (middle), and number of entries to the center (right) after 10 min of exploration. (**B**) Grip strength for forelimbs (left) and hindlimbs (right). (**C**) Pole descent test: latency to return to the home cage (left) and latency to complete T turn on the pole (right). (**D**) Rotarod test for hLRRK2^G2019S^ Tg mice and WT littermate controls at 35 and 50 weeks of age (two-way ANOVA with Sidak; n=8-11). Data are presented as mean ±SEM, \**P*<0.05.

**Fig. S4:**
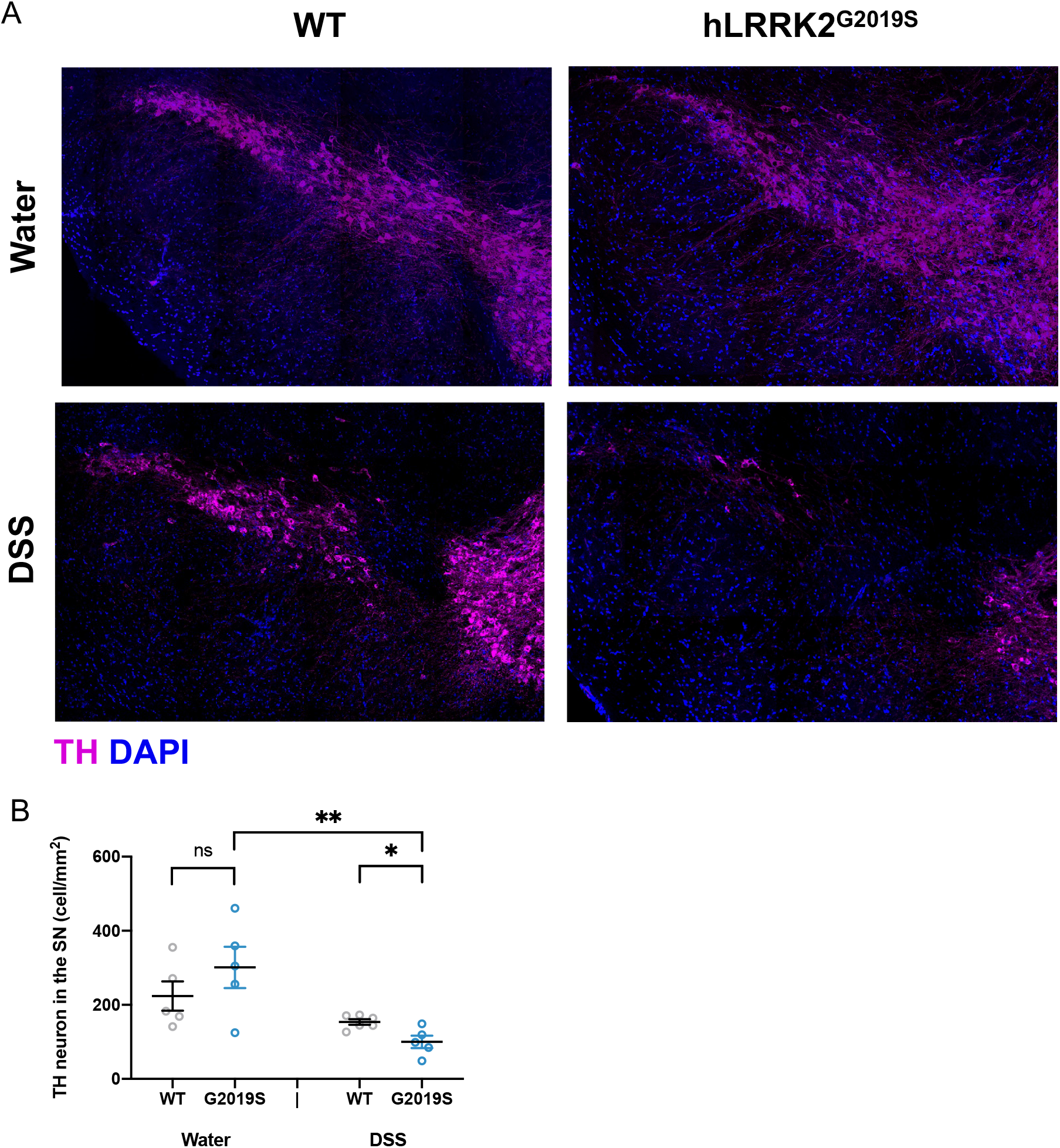
TH neurons in the SN of male hLRRK2^G2019S^ Tg mice after Vehicle (Water) or DSS treatment. (**A**) Representative images of TH^+^ neurons (magenta) in the SN of male hLRRK2^G2019S^ Tg mice and WT littermates at 32 weeks post treatment with DSS or water as a negative control (52 weeks of age). (**B**) Number of TH^+^ neurons in the SN of male hLRRK2^G2019S^ Tg mice and WT littermates at 32 weeks post treatment with DSS or water as a negative control (52 weeks of age) (two-way ANOVA with Sidak; n=5-6). Data for the DSS groups are as in Figure 2G. Data are presented as means ±SEM, \**P*<0.05, \*\*\*\**P*<0.001.

**Fig. S5:**
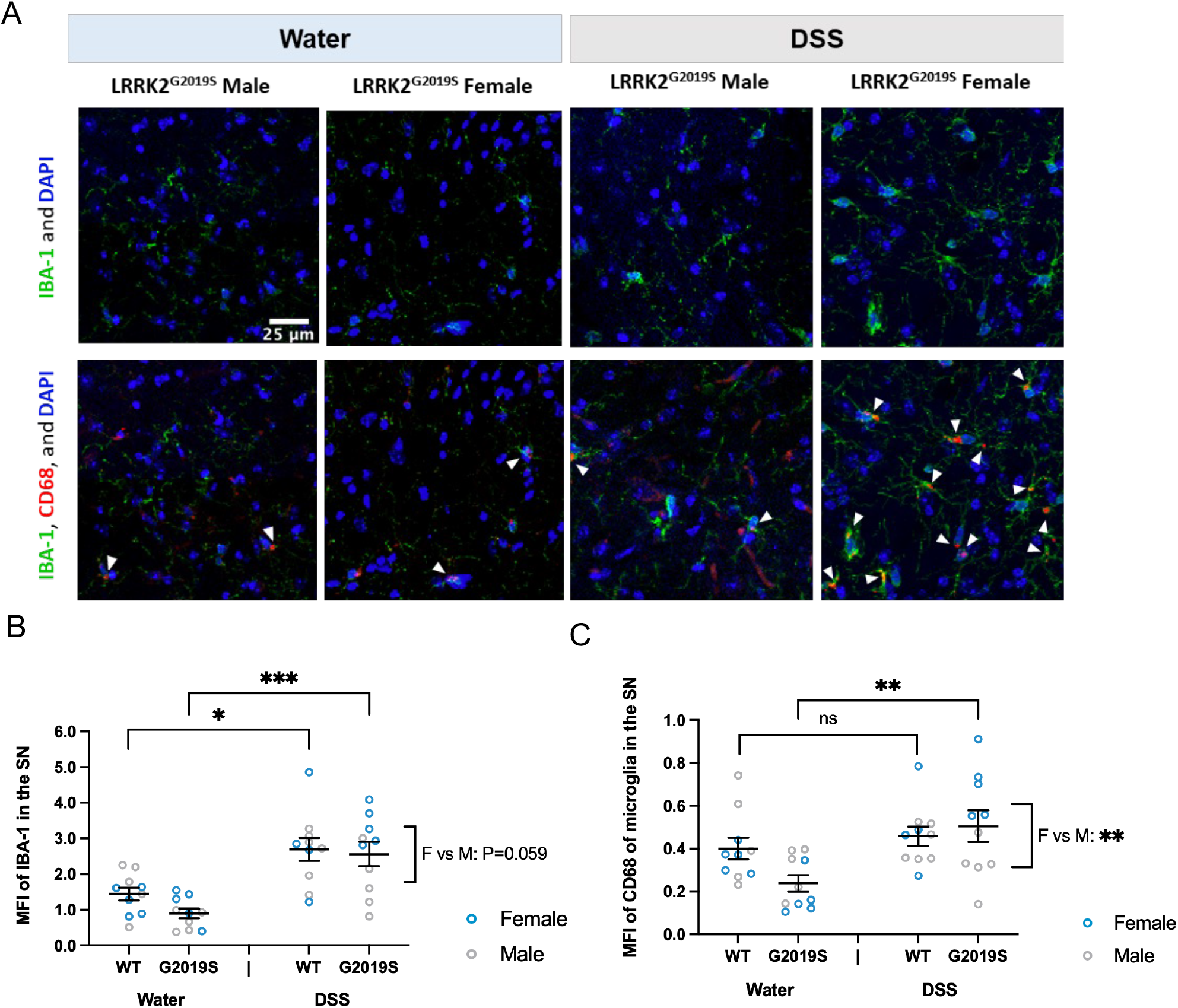
IBA-1 and CD68 in the SN of hLRRK2^G2019S^ Tg mice after DSS treatment. (**A**) Representative images of IBA-1^+^ microglia (green) and CD68 (red) in SN of male and female hLRRK2^G2019S^ Tg mice and WT littermates at 32 weeks post DSS treatment (52 weeks of age). Arrows point to CD68 as puncta within the IBA-1 signal. (**B**) MFI of IBA-1^+^ microglia in the SN. (**C**) MFI of CD68 within the IBA-1^+^ microglia in the SN. (Two-way ANOVA with Sidak; n=4-6). Data are presented as means ±SEM, \**P*<0.05, \*\**P*<0.01, \*\*\**P*<0.001.

**Fig. S6:**
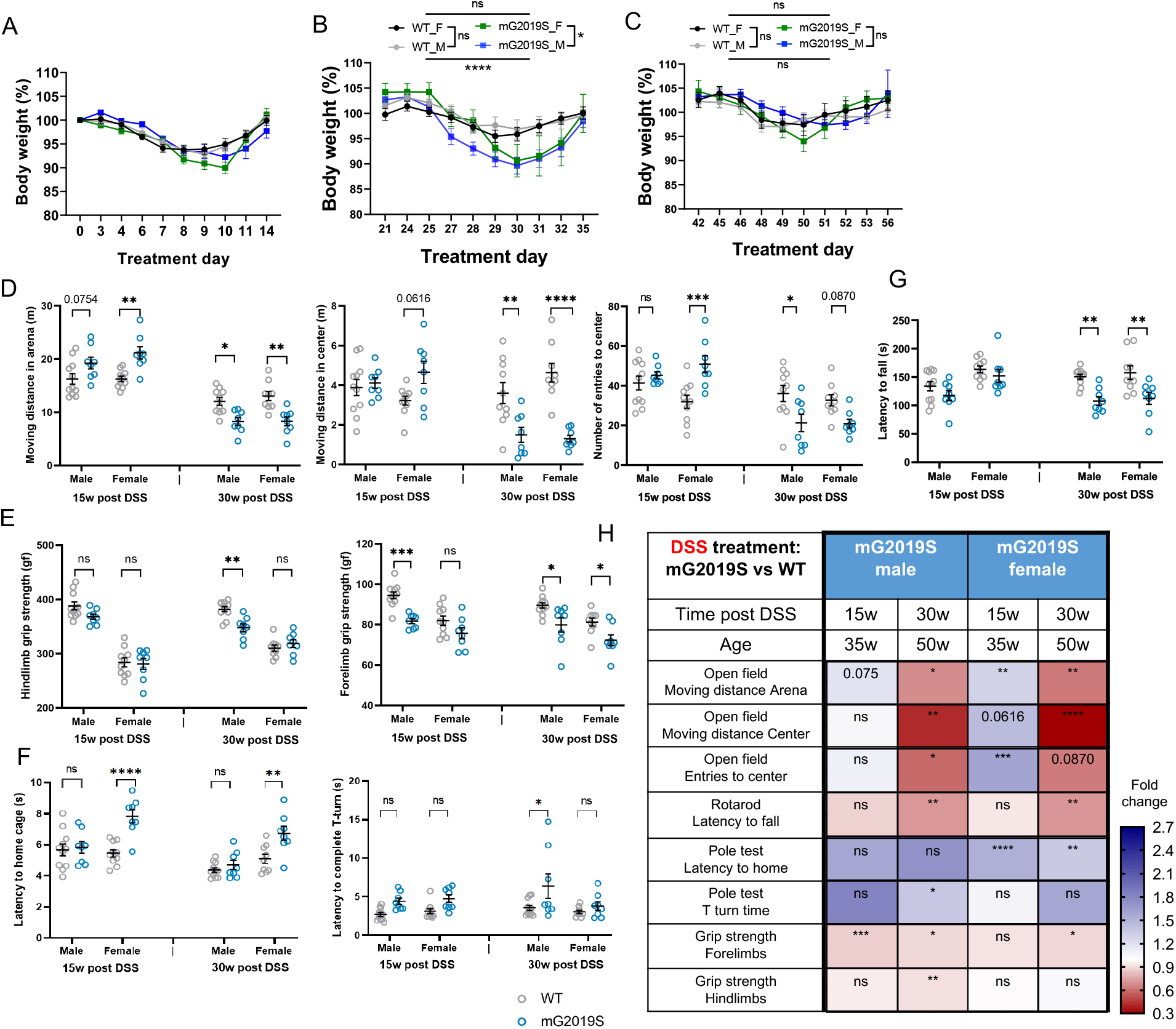
Body weight loss and motor impairments in mLRRK2^G2019S^ KI mice after early DSS treatment. (**A**)-(**C**) Body weight normalized to starting weight (day 0) for male (M) and female (F) mLRRK2^G2019S^ KI mice (mG2019S) and wildtype littermate controls (WT) over 3 rounds DSS treatment (two-way ANOVA with Sidak, n=8-11). (**D**) Open field test: moving distance in the arena (left), center (middle) and number of entries to the center (right) after 10 min exploration, (**E**) Grip strength for the hindlimbs (left) and forelimbs (right), (**F**) Pole descent test: latency to return to the home cage (left) and latency to complete T turn on the pole (right), (**G**) Rotarod test of mLRRK2^G2019S^ KI mice and wildtype littermates at 15 and 30 weeks post DSS treatment. (**H**) Summary of behavioral testing results for mLRRK2^G2019S^ KI mice (mG2019S) relative to WT littermate controls at 15 and 30 weeks post DSS treatment (two-way ANOVA with Sidak, n=8-11). Data are presented as means ±SEM, *P<0.05, \*\**P*<0.01, \*\*\**P*<0.001 and \*\*\*\**P*<0.0001.

**Fig. S7:**
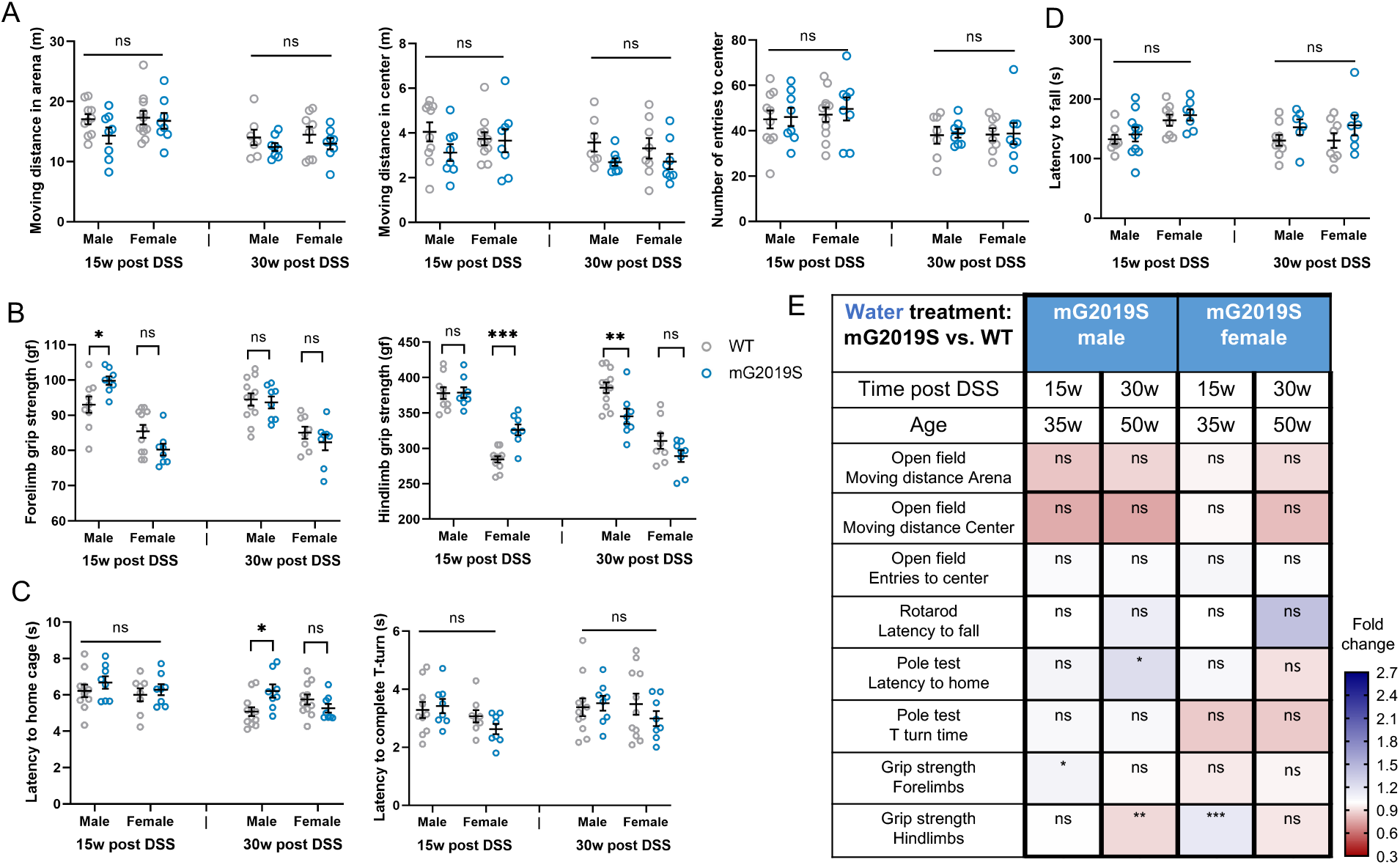
Vehicle-treated (water; untreated) mLRRK2^G2019S^ KI mice and WT littermate controls in behavioral tests for motor ability. (A) Open field test: moving distance in the arena (left), center (middle) and number of entries to the center (right) after 10 min of exploration, (B) Grip strength for the forelimbs (left) and hindlimbs (right), (C) Pole descent test: latency to return to the home cage (left) and latency to complete T turn on the pole (right), (D) Rotarod test for mLRRK2^G2019S^ KI mice (mG2019S) and wildtype (W) littermate controls at 15 and 30 weeks post DSS treatment. (E) Summary of behavioral testing results for mLRRK2^G2019S^ KI mice (mG2019S) and WT littermate controls (two-way ANOVA with Sidak; n= 8-10). Data are presented as means ±SEM, \**P*<0.05, \*\**P*<0.01 and \*\*\**P*<0.001

**Fig. S8:**
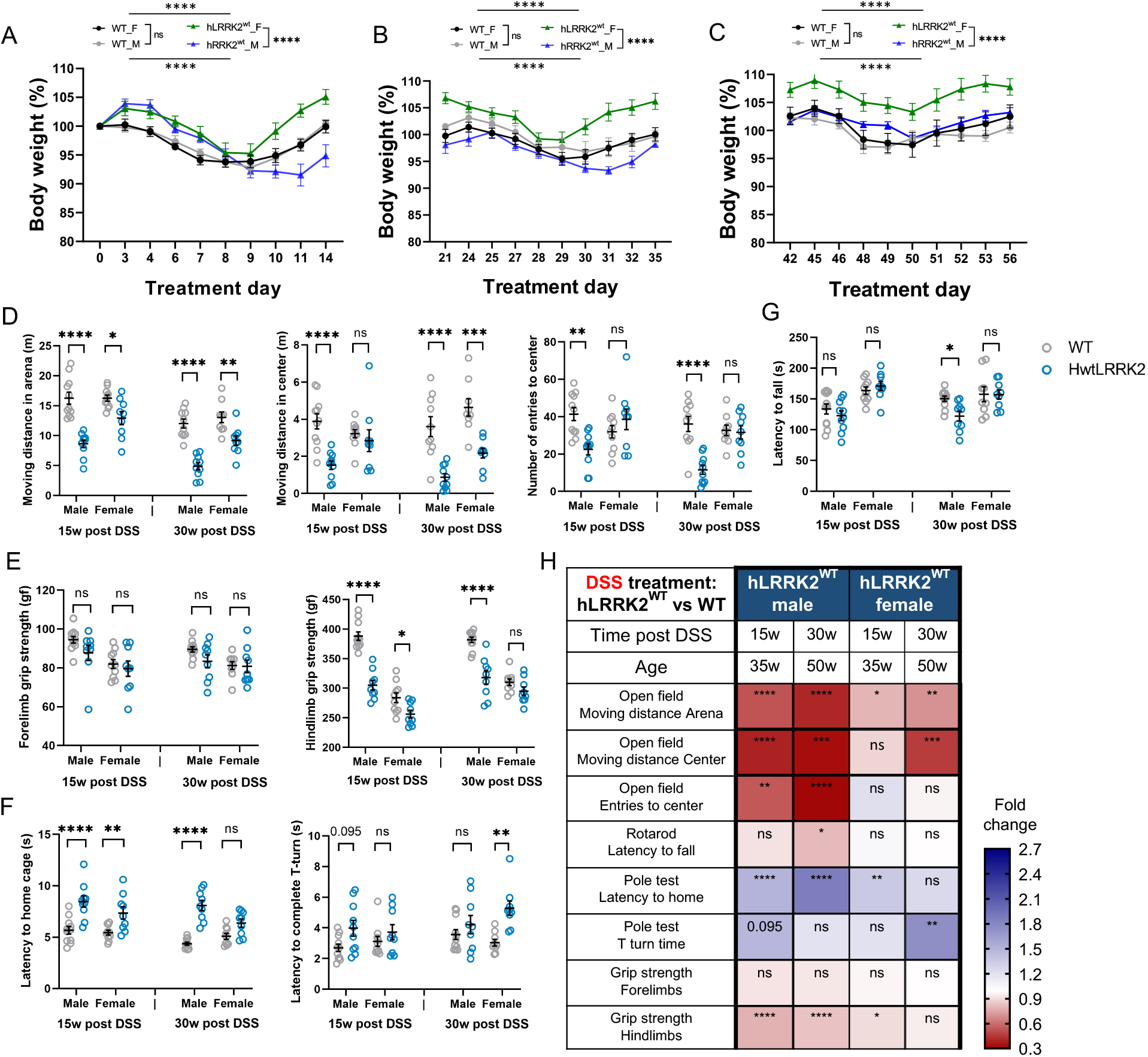
Body weight loss and motor impairments in hLRRK2^WT^ Tg mice after DSS treatment. (**A**)-(**C**) Body weight normalized to starting weight (day 0) for male (M) and female (F) hLRRK2^WT^ Tg mice and wildtype littermates (WT) during each of the 3 rounds of DSS treatment. (**D**) Open field test: moving distance in the arena (left), center (middle) and number of entries to the center (right) after 10 min of exploration, (**E**) Grip strength for the forelimbs (left) and hindlimbs (right), (**F**) Pole descent test: latency to return to the home cage (left) and latency to complete T turn on the pole (right), (**G**) Rotarod test of hLRRK2^WT^ Tg mice and wildtype (WT) littermate controls at 15 and 30 weeks post DSS treatment. (**H**) Summary of behavioral testing results for hLRRK2^WT^ Tg mice and WT controls at 15 and 30 weeks post DSS treatment (two-way ANOVA with Sidak, n=10-11). Data are presented as mean ±SEM, \**P*<0.05, \*\**P*<0.01, \*\*\**P*<0.001 and \*\*\*\**P*<0.0001.

**Fig. S9:**
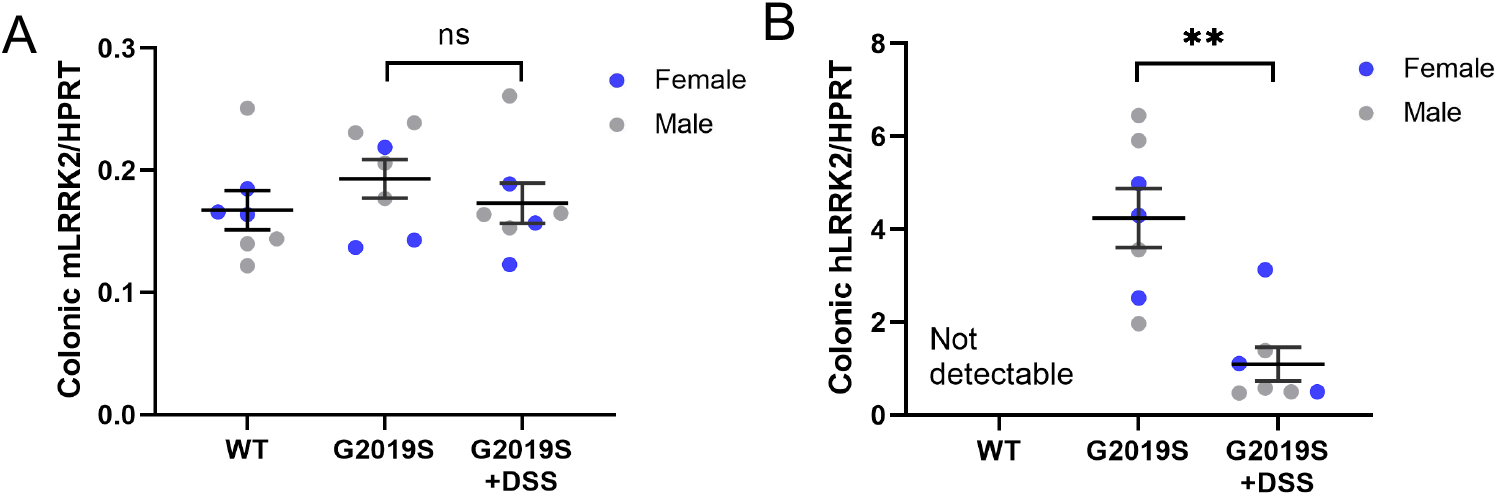
Colonic expression of *LRRK2* in the hLRRK2^G2019S^ Tg mice at baseline and after 7 days of DSS treatment. mRNA levels of endogenous mouse *LRRK2* (**A**) and human *LRRK2 G2019S* transgene (**B**) were examined by real time quantitative PCR in the colon from male and female hLRRK2^G2019S^ Tg mice (G2019S) and wildtype littermates (WT), at baseline and 7 days after DSS treatment (two-way ANOVA with Sidak, n=7). Gene expression is normalized to expression of *HPRT*. Data are presented as mean ±SEM, \*\**P*<0.01.

**Fig. S10:**
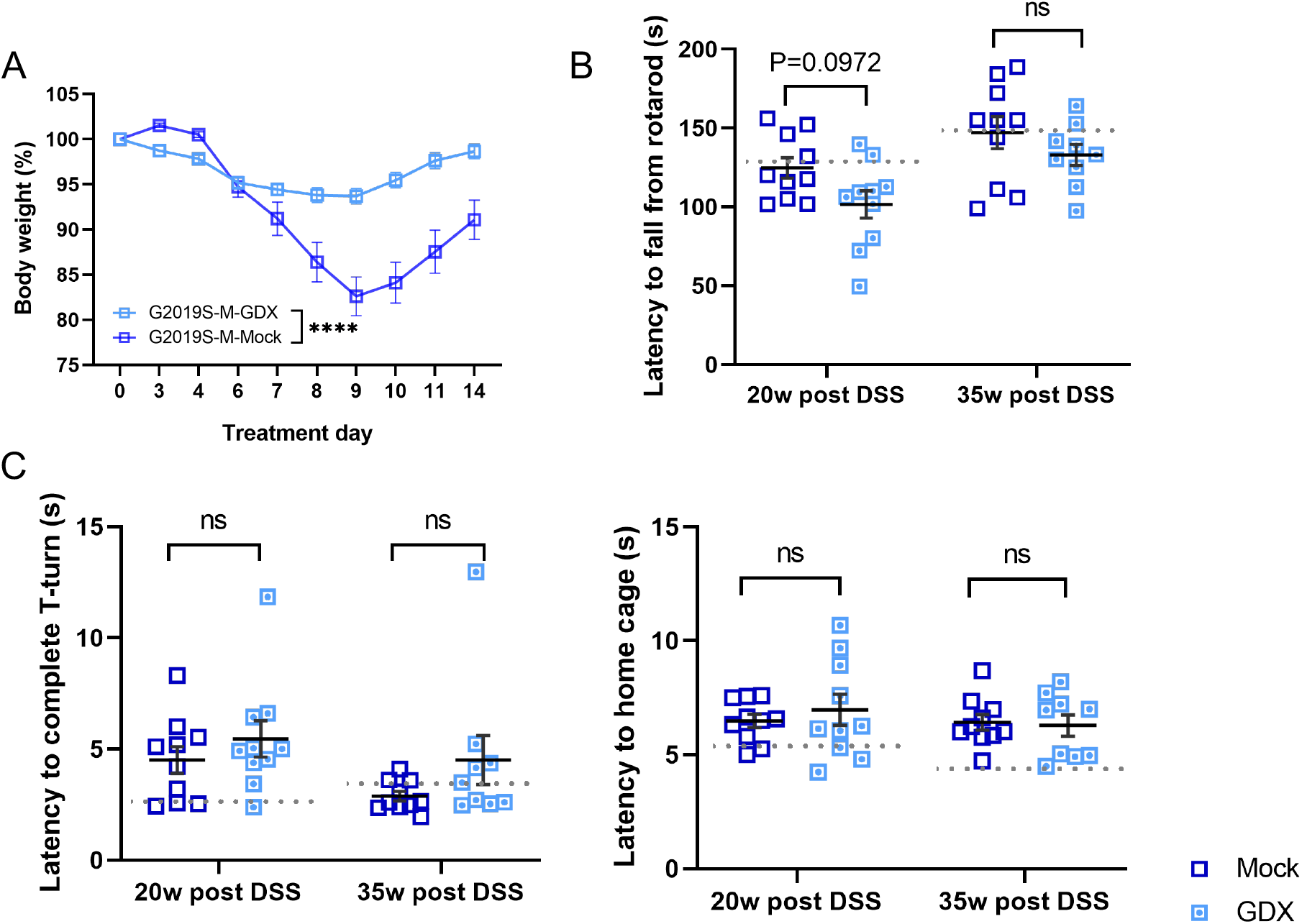
Body weight loss and behavioral performance in additional motor tasks for gonadectomized (GDX) male hLRRK2^G2019S^ Tg mice after DSS treatment. (**A**) Body weight normalized to starting weight (Day 0) for gonadectomized (GDX) or Mock surgery-exposed (Mock) male hLRRK2^G2019S^ Tg mice (G2019S-M) during the first round of DSS treatment. (**B**) Rotarod test and (**C**) Pole descent test: latency to return to the home cage (left) and latency to complete T turn on the pole (right) at 20 and 35 weeks post DSS treatment (two-way ANOVA with Sidak, n=9-10). Data are presented as mean ±SEM, \*\*\*\**P*<0.0001. Dotted lines indicate the mean value for WT littermates as a reference.

**Fig. S11:**
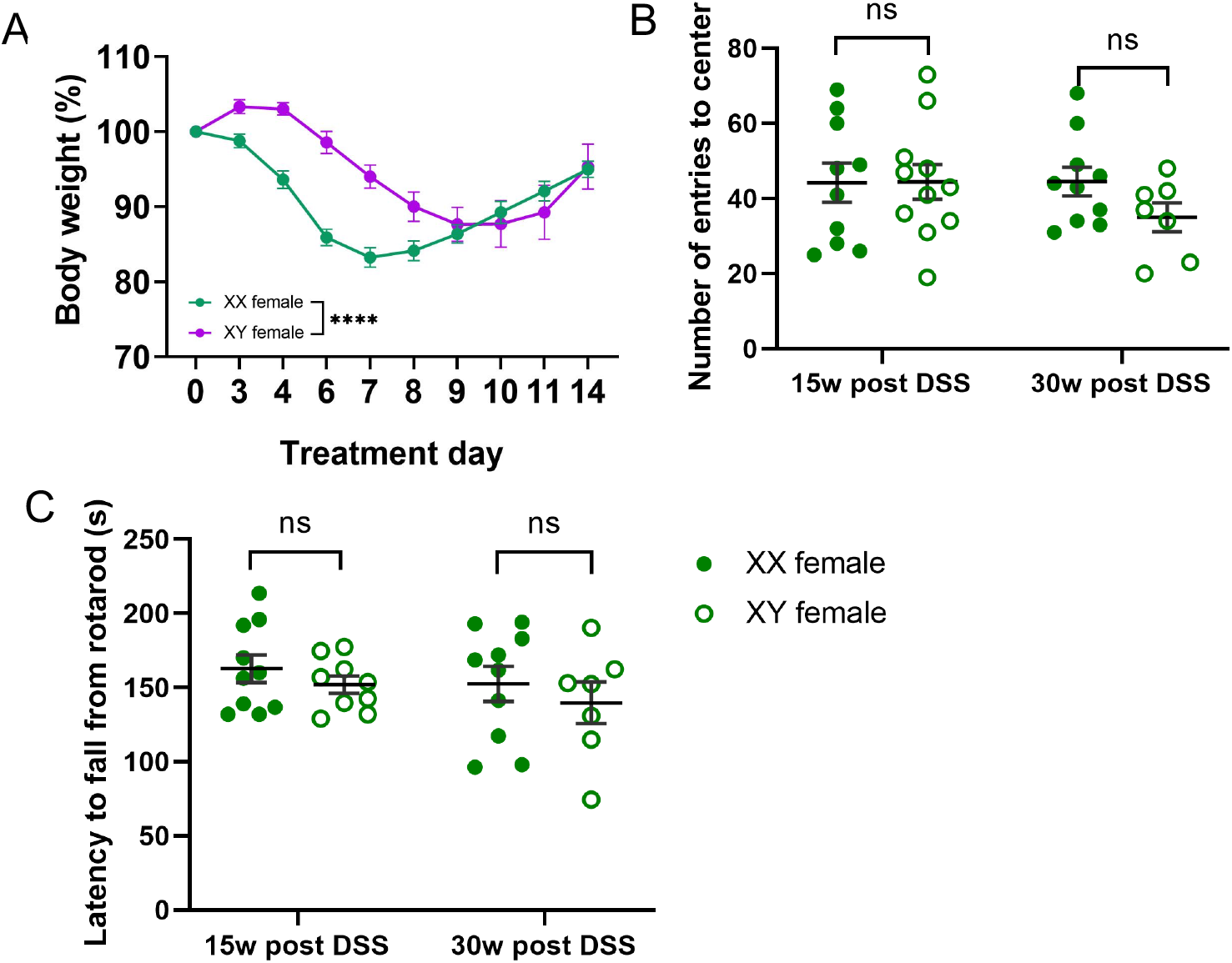
Body weight loss and behavioral performance in additional motor tasks for XY female hLRRK2^G2019S^ Tg mice after DSS treatment. (**A**) Body weight normalized to starting weight (day 0) for XX and XY female hLRRK2^G2019S^ Tg mice during the first round of DSS treatment. (**B**) Open field test: number of entries to the center after 10 min of exploration. (**C**) Rotarod test for XX and XY female hLRRK2^G2019S^ Tg at 15 and 30 weeks post DSS treatment (two-way ANOVA with Sidak, n= 7-10). Data are presented as mean ±SEM, \*\*\*\**P*<0.0001

**Fig. S12:**
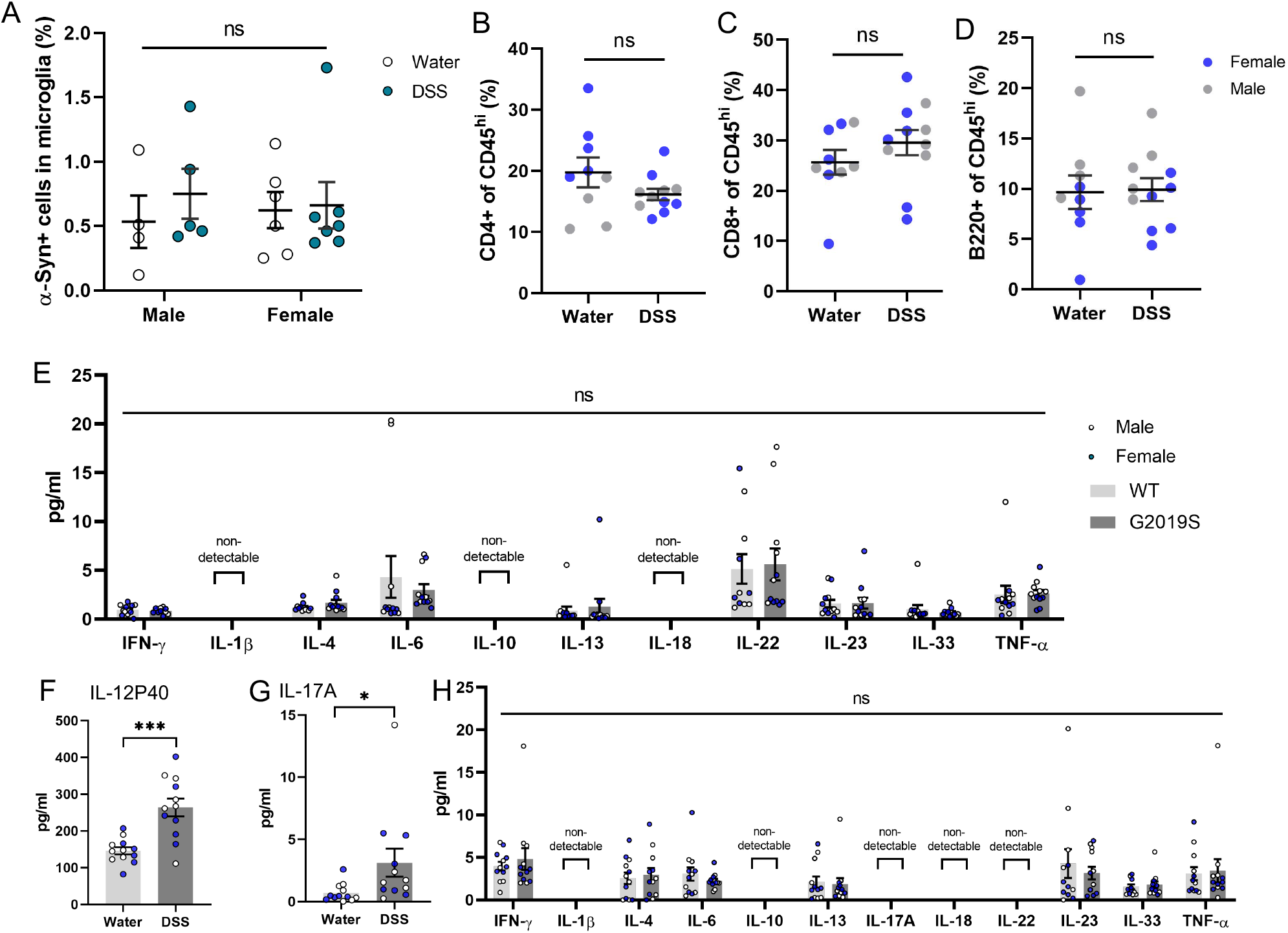
DSS-induced alterations in brain immune profiles of hLRRK2^G2019S^ Tg mice relative to WT littermate controls, prior to the onset of neuropathological and behavioral endophenotypes of PD. Flow cytometry of brain parenchymal cells for (**A**) a-Syn^+^ microglia (two-way ANOVA with Sidak, n=4-6), (**B**) CD4 T cells, (**C**) CD8 T cells and (**D**) B cells from hLRRK2^G2019S^ Tg mice at 1 week after the last round of treatment with DSS or vehicle (water) (paired t-test, n=9-12). (**E**) Serum cytokine concentrations for (**F**) IL-12P40 and (**G**) IL-17A, and (**H**) various cytokines in CSF from hLRRK2^G2019S^ Tg mice at 1 week after the last round of treatment with DSS or vehicle (paired t-test, n=9-12). Data are presented as mean ±SEM, \**P*<0.05 and \*\*\**P*<0.001.

